# Hybridization enables the fixation of selfish queen genotypes in eusocial colonies

**DOI:** 10.1101/2021.01.12.426359

**Authors:** Arthur Weyna, Jonathan Romiguier, Charles Mullon

## Abstract

A eusocial colony typically consists of two main castes: queens that reproduce and sterile workers that help them. This division of labour however is vulnerable to genetic elements that favour the development of their carriers into queens. Several factors, such as intra-colonial relatedness, can modulate the spread of such caste-biasing genotypes. Here we investigate the effects of a notable yet understudied ecological setting: where larvae produced by hybridization develop into sterile workers. Using mathematical modelling, we show that the coevolution of hybridization with caste determination readily triggers an evolutionary arms race between non-hybrid larvae that increasingly develop into queens, and queens that increasingly hybridize to produce workers. Even where hybridization reduces worker function and colony fitness, this race can lead to the loss of developmental plasticity and to genetically hard-wired caste determination. Overall, our results may help understand the repeated evolution towards remarkable reproductive systems (e.g. social hybridogenesis) observed in many ant species.

## 1 Introduction

Eusociality is characterized by a striking division of reproductive labour between two castes: queens and workers (Crespi and Yanega, 1995). Queens monopolize reproduction, while typically sterile workers specialize on other colony tasks such as foraging and tending to the brood. The sterility of workers initially seemed so inconsistent with natural selection that Darwin referred to eusociality as his “one special difficulty” (Darwin, 1859, ch. 7). This apparent paradox was resolved in the 1960’s with W. D. Hamilton’s theory of kin selection (Hamilton, 1964). Hamilton demonstrated that natural selection can favour eusociality when workers preferentially help relatives (who can transmit the same genetic material). In addition to laying the theoretical basis for the evolution of eusociality, Hamilton’s work led to the insight that caste determination should be plastic to allow identical gene copies to be in workers and in the queen they help (Seger, 1981). In line with this notion, the developmental fate of female larvae in many eusocial insects depends on environmental factors (Trible and Kronauer, 2017), such as food quantity and quality (Brian, 1956; Brian, 1973), temperature and seasonality (Brian, 1974; Schwander et al., 2008) or signals emitted by adults of the colony (Penick and Liebig, 2012; Libbrecht et al., 2013). Probably the most iconic example of such plasticity is found in honeybees where queens arise only from larvae reared in royal cells and fed with royal jelly. For long, this and many other empirical findings strengthened the idea that caste determination is under strict environmental control and largely free from genetic effects.

More recently however, substantial genetic variation for caste determination has been described across a number of eusocial species (Winter and Buschinger, 1986; Moritz et al., 2005; Hartfelder et al., 2006; Linksvayer, 2006; Schwander and Keller, 2008; Smith et al., 2008; Frohschammer and Heinze, 2009; Schwander et al., 2010). This variation is thought to derive from caste-biasing genotypes which bias the development of their carrier towards a particular caste (Moritz et al., 2005; Hughes and Boomsma, 2008). Those genotypes that favour larval development towards the reproductive caste have sometimes been referred to as “royal cheats” as they cause the individuals that carry them to increase their own direct reproduction at the expense of other colony members (e.g. Anderson et al., 2008; Hughes and Boomsma, 2008). The segregation of such royal cheats should depend on a balance between: (1) direct benefits from increased representation in the reproductive caste; and (2) indirect costs due to reduced worker production and colony productivity (Hamilton, 1964). As highlighted by abundant theory, several factors can influence these benefits and costs and thus tip the balance for or against the evolution of royal cheats. For instance, low relatedness between larvae due to polyandry (when queens mate with multiple males) or polygyny (when colonies have multiple queens) increases competition between genetic lineages within colonies and thereby favours royal cheating (e.g. Reuter and Keller, 2001). Conversely, selection against cheats is bolstered by low dispersal abilities and high within-group relatedness (e.g. Hamilton, 1964; Lehmann et al., 2008; Boomsma, 2009), bivoltinism and asymetrical sex-ratio (e.g. Trivers and Hare, 1976; Seger, 1983; Alpedrinha et al., 2014; González-Forero, 2015; Quiñones and Pen, 2017), coercion (i.e. policing, Wenseleers et al., 2004; Dobata, 2012), queen longevity and competition between queens (e.g. Queller, 1994; Bourke and Chan, 1999; Avila and Fromhage, 2015), or where workers reproduce following queen death (Field and Toyoizumi, 2020).

One intriguing factor that has been proposed to influence the cost of royal cheating is sperm parasitism, a behavior consisting in queens using the sperm of another species or lineage to produce hybrid workers (Linksvayer, 2006; Anderson et al., 2008). Both morphological and genetic data suggest that this behaviour is common in many ant species (e.g. in multiple *Temnothorax* populations, the majority of queens were found to produce some hybrid workers, Douwes and Stille, 1991; Umphrey, 2006 and Feldhaar et al., 2008 for reviews). In these species, sperm parasitism results in hybrid larvae that rarely, if ever, develop as fertile queens and rather become sterile workers (presumably due to genetic incompatibilities between parental lineages, Feldhaar et al., 2008; Trible and Kronauer, 2017). Such hybrids should therefore be impervious to genetic caste-biasing effects and thus provide a reliable source of workers. In principle, this alternative supply of workers may reduce the indirect cost of royal cheats and hence favours their evolution (Anderson et al., 2008). But be-yond these broad-brush predictions, the effect of sperm parasitism on the segregation of royal cheats remains poorly understood.

Here, we develop a mathematical model to explore the evolution of genetic caste determination via royal cheats when queens can hybridize to produce workers. In particular, we assess the effects of key factors on the evolutionary dynamics of caste determination, such as polyandry and queen parthenogenesis (when queens have the ability to produce daughters asexually), as well as their interactions with potential costs and benefits of hybridization, for instance owing to hybrid incompatibilities or hybrid vigor.

## 2 The model

We consider a large population of annual eusocial haplodiploids with the following life-cycle (fig. 1). First, virgin queens mate with a fixed number *m* ∈ {1, 2, …} of males. Each of these mates can either be an allo-(with probability *η*) or a con-specific male (with complementary probability 1− *η*). Once mated, queens found monogynous colonies (i.e. one queen per colony) and lay a large number of eggs. A proportion *f* of these eggs are diploid (and develop into females) and (1−*f*) are haploid (and develop into males). Assuming random egg fertilization, a queen therefore produces on average *fη* hybrid and *f*(1 − *η*) non-hybrid females. We assume that a hybrid female can only develop as a worker, while a non-hybrid female can either develop as a worker (with probability *ω*) or as a queen (with complementary probability 1− *ω*). Overall, a colony thus consists of *fη* hybrid and *f*(1 − *η*)*ω* non-hybrid sterile workers, as well as *f*(1−*η*)(1−*ω*) virgin queens and (1−*f*) males that are available for reproduction at the next generation.

**Figure 1:**
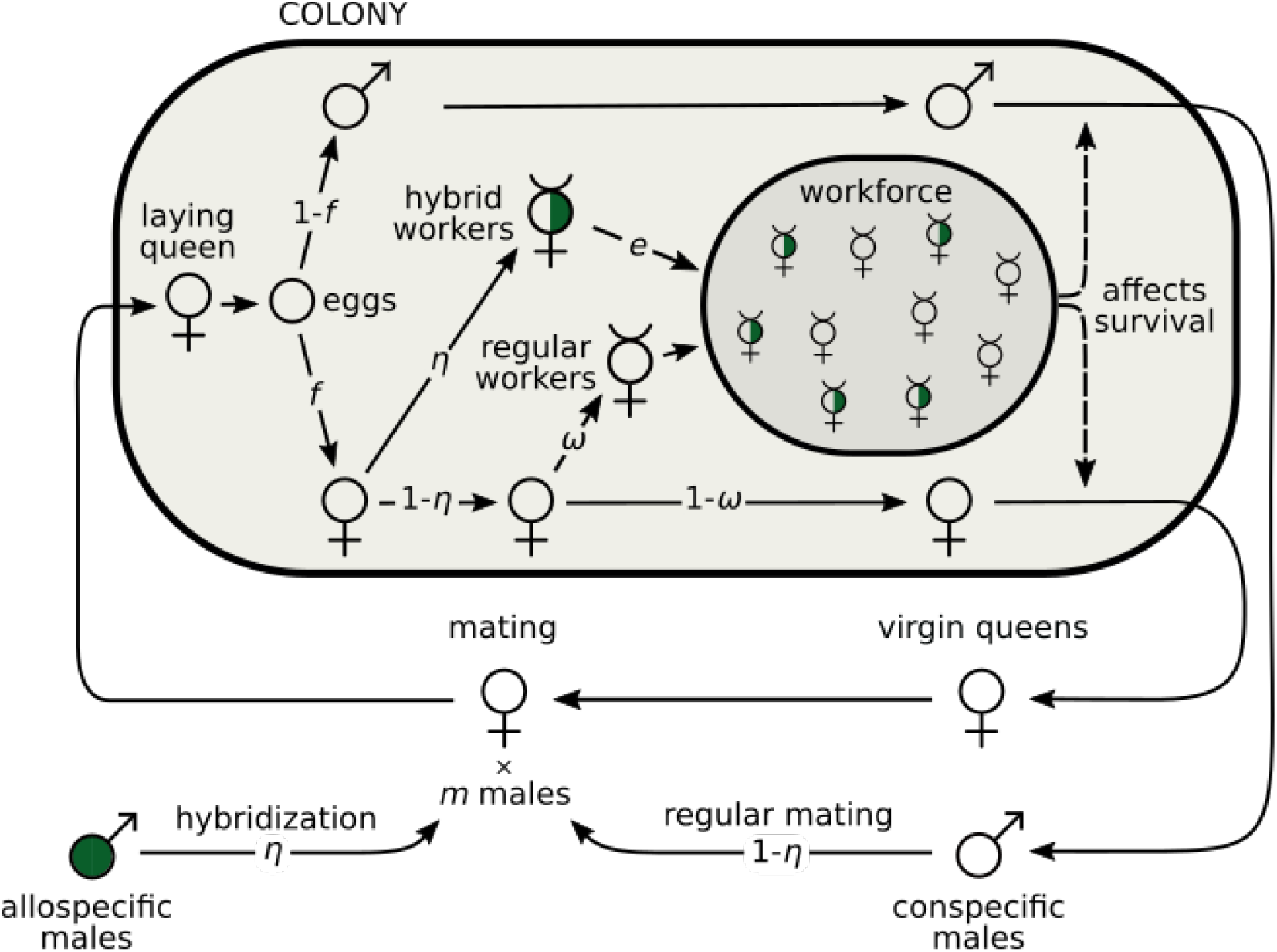
The life cycle of an annual eusocial with hybridization and sperm parasitism. At each generation, the life-cycle begins with virgin queens mating with *m* males, each of which has a probability *η* to be allospecific and 1−*η* to be conspecific. After mating, a queen founds a colony and starts producing eggs. Hybrid female eggs (with allospecific paternal origin) all develop into workers. Regular female eggs (with conspecific paternal origin) develop into workers with probability *ω* and into queens otherwise. The variable *η* thus captures the tendency of queens to hybridize and parasitize sperm, while *ω* controls caste determination.

If only virgin queens and males can reproduce, their reproductive success depends on the workforce of their colony of origin. Specifically, we assume that the probability that a sexual reaches the mating pool increases linearly with the total number of workers in the colony, combining hybrid and nonhybrid workers (we show later that our results do not change qualitatively when the increase is non-linear). We nonetheless allow for differential contribution to the workload between hybrid and non-hybrid workers, with the contribution of hybrid workers weighted by a parameter *e* ≥ 0 (so that the effective workforce of a colony is *efη* + *f*(1 − *η*)*ω*). When *e* = 1, hybrid workers have the same working efficiency as non-hybrid workers. By contrast, when *e* < 1, hybrid workers are less efficient for instance due to outbreeding depression. This can also reflect other potential costs associated with hybridization, such as the production of sterile or non-viable hybrid queens (Feldhaar et al., 2008). Conversely, when *e* > 1 hybrid workers outperform regular workers, due for example to hybrid vigor (Umphrey, 2006).

## 3 Results

### 3.1 Hybridization and sperm parasitism, even costly, can lead to the fixation of royal cheats and the complete loss of intraspecific workers

We first investigate the evolution of caste determination by allowing the probability *ω* that a larva develops as a worker to vary. We assume that this probability is under individual genetic control (i.e. the future caste of a female larva depends only on its own genotype) and that it evolves via random mutations with weak additive phenotypic effects (Appendix A for details on our methods). Mutational effects are unbiased so a new mutation is equally likely to increase or decrease the tendency *ω* of becoming a worker. Those mutations that decrease *ω* can be considered as more selfish as they increase the likelihood that their carriers develop into queens at the expense of other individuals of the same colony. Following the terminology of Hughes and Boomsma (2008), we thus refer to mutations decreasing *ω* as “royal cheats”. As a baseline, we consider the case where queens mate with a large number of males (i.e. *m* → ∞) and where hybridization is fixed at a given level (e.g. *η* is the proportion of allo-specific males in the pool of mates from which females choose randomly).

Our analyses (Appendix B.1.1) reveal that the probability for a larva to develop as a worker evolves towards a unique and stable equilibrium,

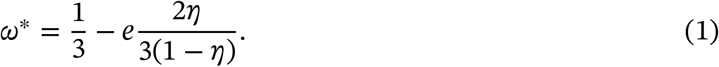

To interpret this equation (1), consider first the case where hybridization is costless (*e* = 1). Eq. (1) then tells that in the absence of hybridization (*η* = 0), a larva will develop into a worker with a probability of 1/3 at equilibrium (in line with previous models that ignore hybridization, e.g. Reuter and Keller, 2001, Appendix B.1.4 for connection). But as hybridization increases (*η* > 0), royal cheating is increasingly favored and larvae become increasingly likely to develop as queens rather than workers (i.e. *ω*^∗^ < 1/3, fig. 2A). In fact past a threshold of hybridization (*η* ≥ 1/3), the population evolves towards a complete loss of non-hybrid workers via the fixation of increasingly caste-biasing royal cheats alleles (*ω* → 0). In this case, non-hybrid females eventually all develop into queens that rely on sperm parasitism to produce workers.

**Figure 2:**
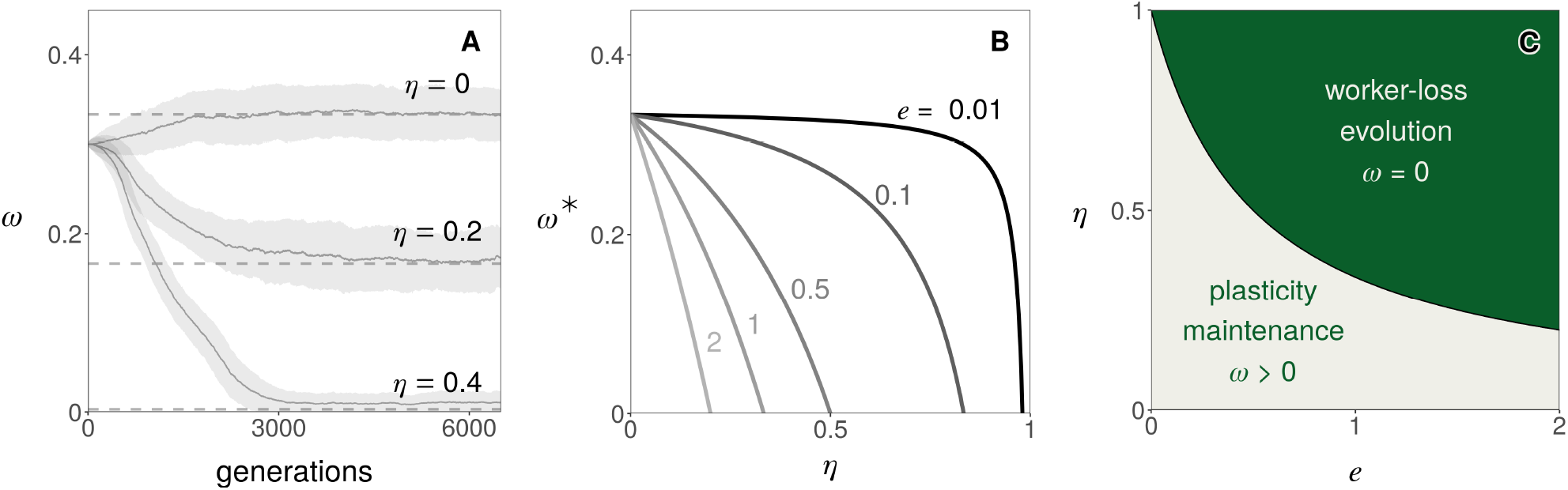
The fixation of royal cheats and evolution of intraspecific worker-loss. **A** Evolution of the probability *ω* that a female larva develops into a worker in a simulated population when queens mate with a large number of males (polyandry, *m* → ∞) and the proportion of allospecific males *η* is fixed (top *η* = 0; middle *η* = 0.2, bottom *η* = 0.4; other parameters: *e* = 1, Appendix A.3 for details on simulations). Plain lines (and surrounding grey areas) show the population average *ω* (and its standard deviation). Dashed lines show the predicted equilibrium (from eq. 1). **B** Equilibrium of *ω* as a function of hybridization *η* and the efficiency of hybrid workers *e* (from eq. 1). **C** Parameter combinations leading to the evolution of complete worker-loss (i.e. *ω* → 0, in green, corresponding to *η* ≥ 1/(1+2*e*) which is found by substituting eq. 1 into *ω*^∗^ ≤ 0).

Equation (1) also shows that the performance of hybrid workers relative to non-hybrids, *e*, modulates the effect of hybridization on the evolution of caste determination (fig. 2B). As a result, royal cheating and worker-loss evolution are facilitated when hybrids outperform regular workers (*e* > 1) but hindered otherwise (*e* < 1). Nevertheless, even where hybridization is extremely costly (0 < *e* ≪ 1), there exists a threshold of hybridization above which complete worker-loss evolves (fig. 2C).

### 3.2 Worker-loss readily emerges from the coevolution of genetic caste determination and sperm parasitism, driven by intra-colonial conflict

The above analysis indicates that intraspecific worker-loss can evolve when queens have a sufficiently high tendency to hybridize. This raises the question of whether such tendency is also subject to selection. To answer this question, we allow the probability *η* that a queen’s mate is allospecific to coevolve with caste determination (*ω*). We assume that this probability *η* is under individual queen control (i.e. it depends only on a queen’s genotype) and like caste determination, evolves via rare mutations with weak additive phenotypic effects (Appendix A for details).

We find that depending on the efficiency *e* of hybrid workers, the coupled evolutionary dynamics of hybridization *η* and caste determination *ω* lead to an evolutionary arms race with one of two contrasted outcomes (Appendix B.1.2 for analysis). When *e* is small (*e* ≤ 1/4, fig. 3A gray region), the population evolves hybridization avoidance (*η* → 0) while the probability *ω* to develop as a worker stabilises for its baseline equilibrium (*ω*^∗^ = 1/3, fig. 3B). By contrast, when hybrid workers are at least half as efficient as regular workers (*e* ≥ 1/2, fig. 3A, dark green region), intraspecific worker-loss evolves (*ω* → 0) and hybridization stabilizes at an intermediate equilibrium (*η*^∗^ = 2/3, fig. 3D). When hybrid worker efficiency is intermediate (1/4 < *e* < 1/2, fig. 3A, light green region), the population evolves either hybridization avoidance or intraspecific worker-loss depending on initial conditions (fig. 3C), with worker-loss favoured by high initial tendency *η* of queens to hybridize. In sum, provided four hybrid workers are at least as good as one regular worker (*e* > 1/4), the coevolution of genetic caste determination and hybridization can lead to worker-loss in our model.

**Figure 3:**
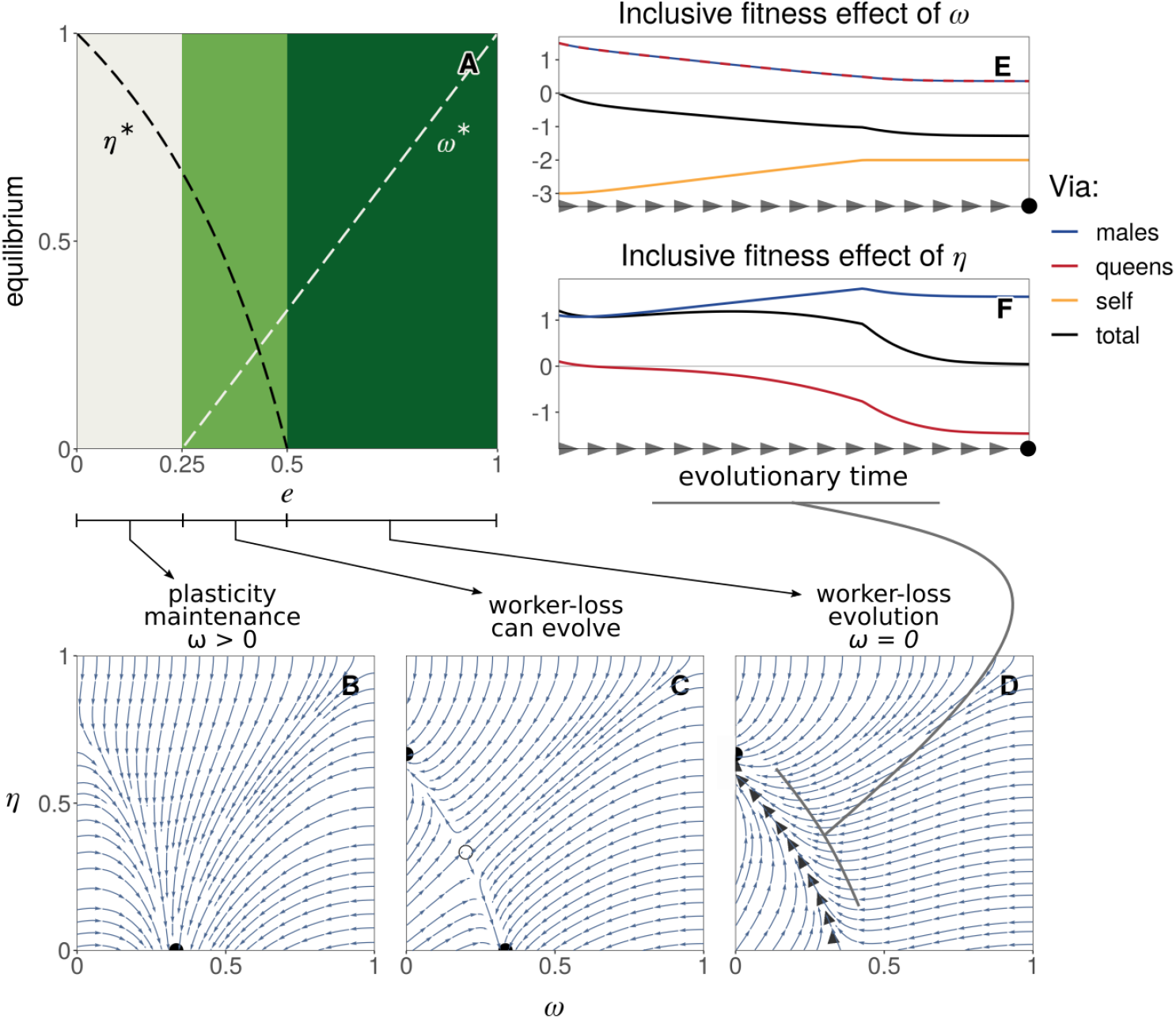
The coevolution of caste determination and sperm parasitism. **A**. Evolutionary equilibria (for *η* in black and *ω* in white) as a function of hybrid worker efficiency *e* (eq. B-6 in Appendix B.1.2 for details). These equilibria however are evolutionary repellors (eq. B-7 in Appendix B.1.2). As a result, three types of coevolutionary dynamics are possible depending on *e* as illustrated in panels B, C & D (from eq. B-5). These panels show examples of phenotypic trajectories when worker-loss: **B** never evolves (*e* = 0.1); **C** can evolve depending on initial conditions (*e* = 0.4); **D** always evolves (*e* = 0.7). Black filled circles indicate the two evolutionary end-points: hybridization avoidance with developmental plasticity (*ω* = 1/3 and *η* = 0 in B-C) or worker-loss with hybridization (*ω* = 0 and *η* = 2/3 in C-D). Empty circle in C shows the internal unstable equilibrium (eq. B-6). Thick grey arrow heads in D represent the trajectory of a population starting from *ω* = 1/3 and *η* = 0 and evolving to worker-loss. **E**: Fitness effects of caste determination *ω* in a mutant larva via itself (in orange), related queens (red) and related males (blue) along the trajectory leading to worker-loss shown in panel D (total selection in black, Appendix B.1.3 for derivation). We see that negative fitness effects via self (orange line) lead to a total selection effect that is negative (black line). This indicates that mutant larvae with increasingly small values of *ω* are selected because these values increase larvae’s direct fitness (by increasing the probability that they develop into queens). **F**: Fitness effects of hybridization *η* in a mutant queen, via its sons (blue) and daughter queens (red) along the trajectory leading to worker-loss shown in panel D (total selection in black). Positive total selection (in black) is mostly due to an increase of fitness via males (in blue). This says that mutant queens with increasingly large values of *η* are selected because this increases their reproduction, especially via males.

To better understand the forces at play in the emergence of worker-loss, we further used a kin-selection approach to decompose the invasion fitness of mutant alleles into the sum of: (1) their direct fitness effects on the reproductive success of the individuals that express them; and (2) of their indirect fitness effects on other related individuals that can also transmit them (Taylor and Frank, 1996, Appendix B.1.3 for details). Starting with a population at the baseline equilibrium in absence of hybridization (*ω* = 1/3, *η* = 0), we tracked these different fitness effects along a typical evolutionary trajectory that leads to worker-loss (black arrow heads, fig. 3D) for alleles that influence the tendency of a larva to develop as a worker (fig. 3E) and of a queen to hybridize (fig. 3F).

Our kin selection analysis reveals that alleles which increase hybridization in queens are selected because they allow queens to increase the number of sexuals produced by their colony (especially via males, blue curve, fig. 3F). This is because the baseline tendency *ω* to develop as a worker that evolves is optimal from the point of view of a gene in a larvae, but sub-optimal from the point of view of a gene in a queen who would benefit from a larger workforce. Hybridization by queens evolves to rectify this and align colony composition with the interests of the queen. Simultaneously, as queens evolve greater hybridization and augment their workforce with hybrids, genes in non-hybrid larva have an increasing interest for their carriers to develop as queens rather than workers (fig. 3E). These two selective processes via queens and larvae fuel one another in an evolutionary arms race whose endpoint is complete intraspecific worker-loss. Our decomposition of fitness effects thus shows that the loss of non-hybrid workers evolves in our model due to within-colony conflicts over colony composition. In fact, our results suggests that worker-loss emerges because hybridization allows queens to control the production of workers in their colony, while non-hybrid larvae lose their tendency to develop as workers to promote their own reproduction via the fixation of royal cheats.

### 3.3 Worker-loss is impaired by low polyandry but facilitated by asexual reproduction

So far, we have assumed that queens mate with a large, effectively infinite, number of males. By increasing relatedness within the brood, low polyandry (2 ≤ *m* ≪ ∞) and monandry (*m* = 1) mediate within-colony conflicts and therefore should be relevant to the evolutionary arms race leading to worker-loss (Anderson et al., 2008; Schwander et al., 2010). To test this, we investigated the effect of mate number *m* on the coevolution of *ω* and *η* (Appendix B.2.1 for details).

We find that as the number *m* of mates decreases, the conditions for intraspecific worker-loss emergence become more restrictive. Specifically, the threshold of hybrid worker efficiency *e* above which worker-loss always evolves increases as polyandry decreases (as *m* → 1, fig. 4A, dark green region). In addition, when the number of mates is low (*m* ≤ 4), evolutionary dynamics do not necessarily lead to either complete worker-loss or hybridization avoidance. For intermediate values of *e* (fig. 4A, blue region) the population actually converges to an intermediate state where queens partially hybridize (0 < *η*^∗^ < 1) and larvae retain developmental plasticity (0 < *ω*^∗^ < 1, fig. 4B, Appendix B.2.1 and fig. S1 for analysis). Under monandry (*m* = 1) the evolution towards such intermediate state always happens when hybrid workers outperform regular workers (*e* > 1, fig. 4A, blue region).

**Figure 4:**
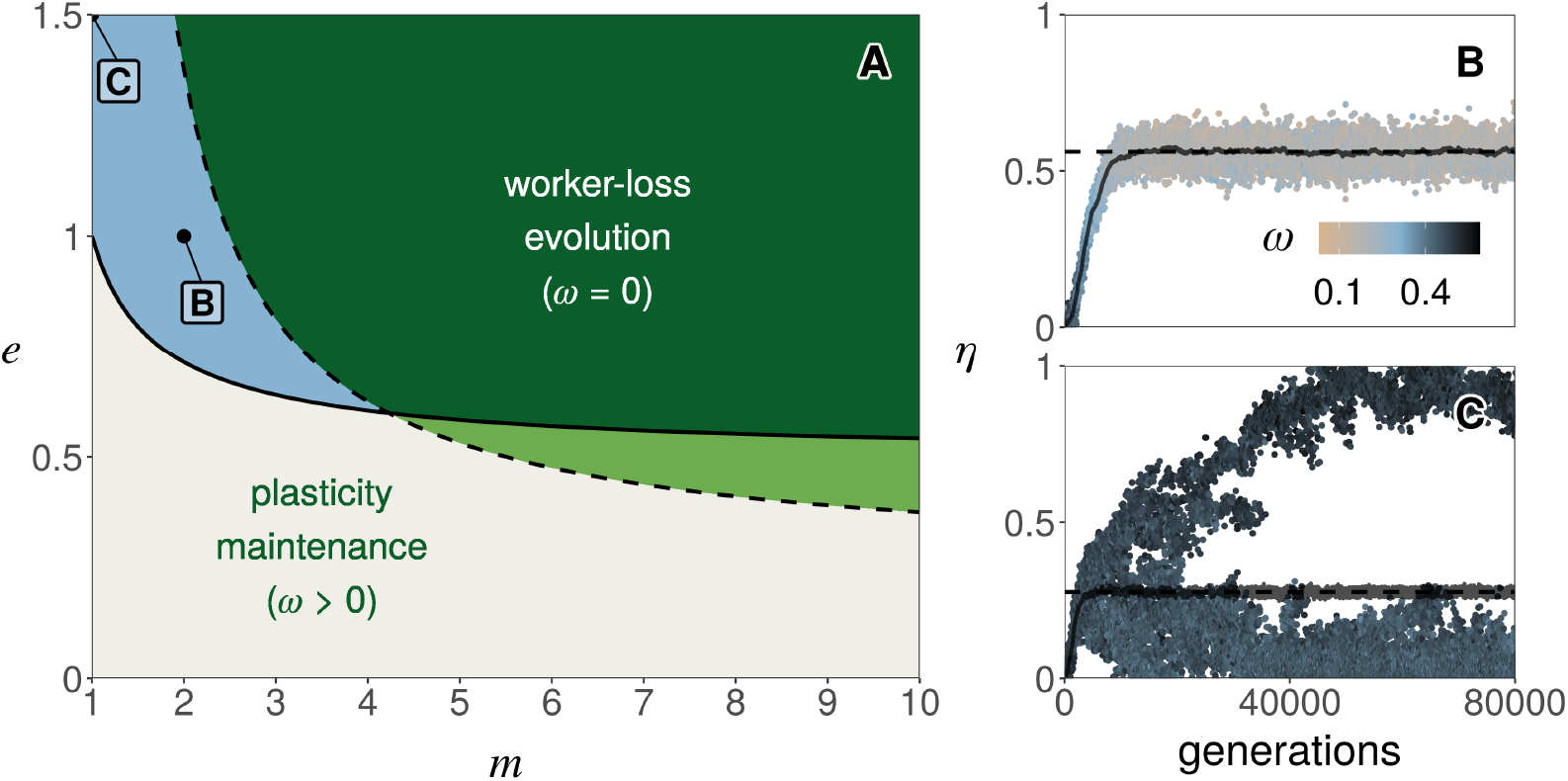
The effects of monandry and low polyandry. **A**: Outcome of selection as a function of mate number *m* and hybrid worker efficiency *e*. Over the dashed line, worker-loss is a stable equilibrium (i.e. a population with traits *ω* = 0 and *η* = 2/3 cannot be invaded, eq. B-16 in Appendix B.2.1). Over the plain line, hybridization can invade when rare (i.e. *η* = 0 is unstable, eq. B-18 in Appendix B.2.1). Below both lines (gray region), plasticity in caste determination is maintained (as in fig. 3B). Over both lines (dark green region), hybridization and worker-loss evolve (as in fig. 3D). In the light green region, worker-loss evolve for some initial conditions (as in fig. 3C). In the blue region, there exists an internal attractor equilibrium (i.e. the population converges towards a phenotype 0 < *η*^∗^ < 1 and 0 < *ω*^∗^ < 1) that is either uninvadable (for 2 ≤ *m* ≤ 4, panel B for e.g.) or invadable leading to polymorphism (for *m* = 1, panel C for e.g.). **B**: Evolution towards an uninvadable phenotype in a simulated population (when *e* = 1 and *m* = 2). Each dot represents the value of *η* of one of 20 haplotypes randomly sampled every 100 generation in a simulated population of 10000 queens (Appendix A.3 for details on simulations). The colour of each dot gives the value of *ω* of the associated haplotype (legend). The horizontal dashed line represents the predicted equilibrium (from fig. S1). The grey line represents the mean value of *η* across the simulation. **C**: Evolution towards an invadable phenotype and the emergence of polymorphism in a simulated population (when *e* = 1.5 and *m* = 1, other parameters and figure legend: same as B).

In the special case of monandry and overperforming hybrid workers (*m* = 1 and *e* > 1), our mathematical analysis further shows that partial hybridization and larval plasticity is not evolutionary stable (Appendix B.2.1, figs. S1-S2). Rather, the population experiences disruptive selection which should favour the emergence of polymorphism. To test this, we performed individual based simulations under conditions predicted to lead to polymorphism (fig. 4C). These show the emergence and long-term coexistence of two types of queens: one which hybridizes with low probability (and reproduces via both males and queens); and another which mates almost exclusively with allospecific males and thus reproduces mostly via males (because *m* = 1, these queens only produce hybrid workers and males). Beyond this special case, the evolution of worker-loss is impeded by low polyandry and impossible under monandry in our model. This is because with a low number of mates, a queen runs the risk of being fertilized by only one type of male. Under complete worker-loss (when the population is fixed for *ω* = 0), a queen mated to only conspecific males produces only larvae destined to be queens but no workers to ensure their survival and thus has zero fitness.

Our finding that monandry inhibits the emergence of worker-loss contrasts with the observation that several ant species, notably of the genus *Cataglyphis*, lack non-hybrid workers and rely on sperm parasitism for workers in spite of being mostly monandrous (Kuhn et al., 2020). One potential mechanism that could have allowed such evolution is thelytokous parthenogenetic reproduction by queens, whereby queens can produce daughters clonally. This reproduction mode, which is common in eusocial Hymenoptera (Rabeling and Kronauer, 2013) and in particular in *Cataglyphis* (Kuhn et al., 2020), could allow queens fertilized exclusively by allospecific males to nevertheless produce queens via parthenogenesis. To investigate how thelytokous parthenogenesis influences the evolution of caste determination, we extend our model so that a fraction *c* of the female progeny of queens is produced parthenogenetically (Appendix B.2.2 for details). We assume that larvae produced in such a way are equivalent to non-hybrid larvae: they develop into workers with a probability *ω* determined by their own genotype (which in this case is the same as their mother’s genotype) and if they develop into workers, they have the same working efficiency as non-hybrid workers (i.e. there is no direct cost or benefit to parthenogenesis).

The coevolutionary dynamics of caste determination and hybridization with parthenogenesis are in general too complicated to be tractable. We could nonetheless gain insights into worker-loss evolution by performing an invasion analysis, asking (1) when is worker-loss (*ω* = 0) evolutionary stable (so that a population where intraspecific workers have been lost cannot be invaded by a genetic mutant with developmental plasticity)? And (2) when can hybridization evolve when absent in the population (i.e. when is *η* = 0 evolutionary unstable)? When these two conditions are met, evolution will tend to favour the emergence and maintenance of worker-loss (as in fig. 3D for e.g.). We thus studied when conditions (1) and (2) above are both true in terms of parthenogenesis *c*, as well as hybrid workers efficiency *e* and mate number *m*. This revealed that parthenogenesis has a non-monotonic relationship with worker-loss evolution (fig. 5A & 5B). As parthenogenesis increases from zero, worker-loss evolution is initially favoured, especially under monandry (as expected; fig. 5C for e.g.; see eq. (B-26) in the Appendix for details). But past a threshold of parthenogenesis, the conditions leading to worker-loss become increasingly stringent until such evolution becomes impossible (see eq. (B-25) in the Appendix for details). This is because as parthenogenesis increases, the relatedness among a queen and larvae of the same colony also increases. The conflict between them, which fuels the evolution of worker-loss, therefore abates until it is no longer advantageous for a larva to preferentially develop as a queen.

**Figure 5:**
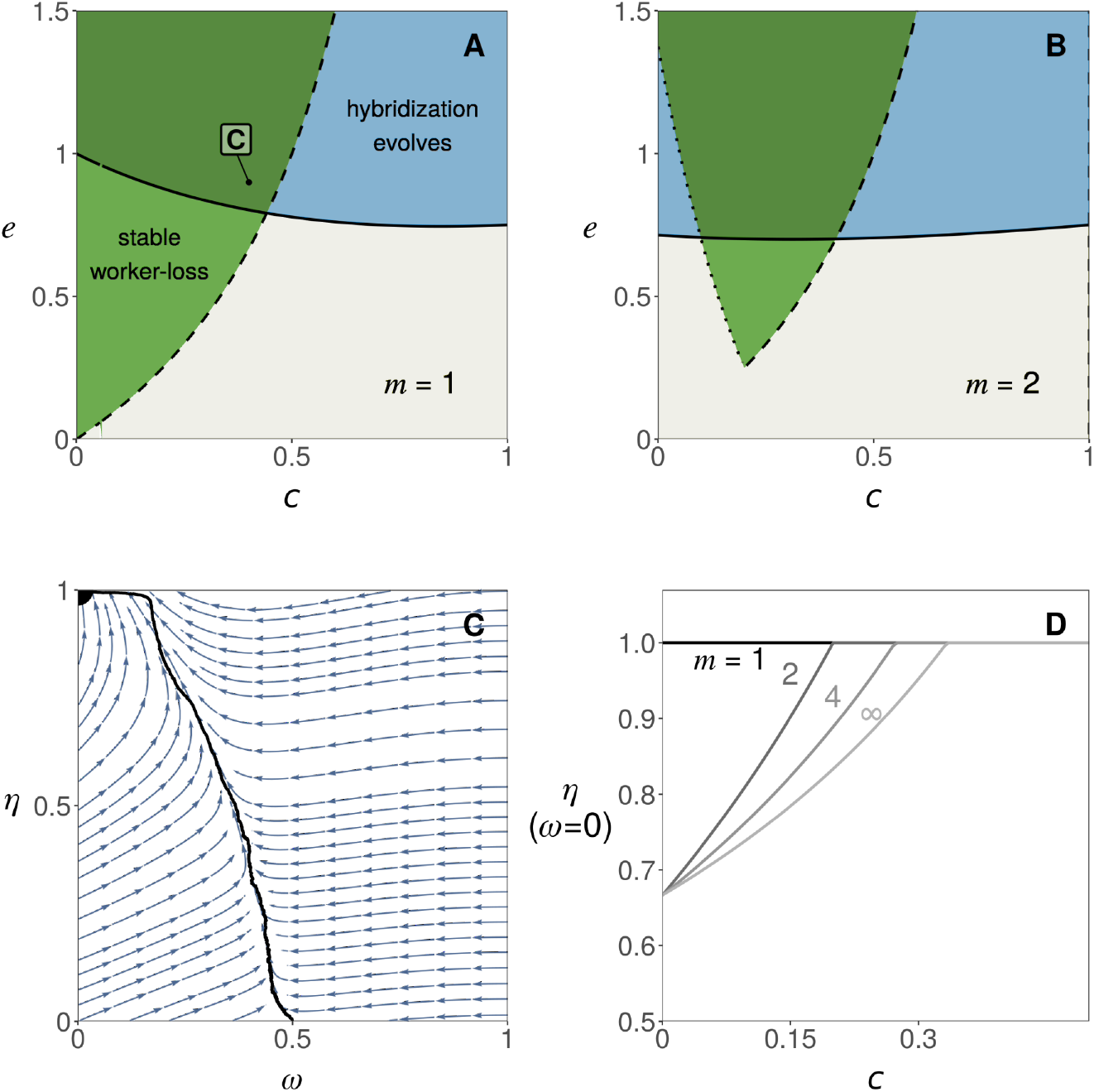
The influence of thelytokous parthenogenesis. **A & B** Invasion analysis as a function of parthenogenesis *c* and hybrid worker efficiency *e* (with *m* = 1 in A and *m* = 2 in B). In the region over the plain line, hybridization can invade when rare (i.e. *η* = 0 is unstable, eq. B-23). In the region over the dashed line (in **A**) or framed by the dotted and dashed lines (in **B**), worker-loss is a stable equilibrium (i.e. a population at equilibrium for *η* and with *ω* = 0 cannot be invaded, Appendix B.2.2, eqs. B-25 and B-26 for details). In the dark green region, selection thus favours both the evolution of hybridization and the maintenance of worker-loss (e.g. panel C). In the light green region, worker-loss can evolve only for some initial conditions (as in fig. 3C). **C** Phenotypic trajectories leading to worker-loss (when *e* = 0.9, *c* = 0.4 and *m* = 1). Arrows show the direction of evolution favoured by selection. Black filled circles indicate the evolutionary end-point. The black line shows the average trait values of a simulated population starting at (*ω* = 1/2, *η* = 0). In this example, selection leads to a state where worker-loss (*ω* = 0) is coupled with complete hybridization (*η* = 1). **D** Level of hybridization *η* favoured by selection when worker-loss has evolved (*ω* = 0) as a function of parthenogenesis *c*. This shows that worker-loss is always associated to complete hybridization (*η* = 1) under monandry (*m* = 1) and if *c* ≥ (*m* − 1)/(3*m* − 1) under polyandry (*m* > 1) (Appendix B.2.2, eq. B-24, for details).

We additionally computed the level of hybridization favoured by selection when the population has evolved worker-loss (and this is an evolutionarily stable state). We find that hybridization increases as queens mate with fewer males and as parthenogenesis increases (fig. 5D), so much so that selection can lead to complete hybridization (*η* = 1, e.g. fig. 5C). As a result, there exists a range of intermediate values of parthenogenesis for which worker-loss evolves in association with a complete loss of intraspecific matings, i.e. queens never mate with males of their own species or lineage. These males are nevertheless still being produced in our model (as the primary sex ratio is such that *f* < 1).

## 4 Discussion

In sum, our analyses indicate that worker-loss readily evolves when queens can hybridize with a lineage of males by whom fertilization leads to the production of workers. This evolution in our model occurs through a sequence of substitutions of alleles that increasingly bias the development of their carrier towards the queen caste, i.e. “royal cheats”. Hybridization, or sperm parasitism, allows royal cheats to fix in the population by providing a way for colonies to compensate for the reduced workforce. In fact, when queens are capable of recognising genetic differences among males and when royal cheats are present in the population, selection favours hybridization by queens to regain control over caste allocation in their colony. This is turn promotes greater cheating by larvae, which favours greater hybridization by queens and so on. This evolutionary arms race, fuelled by intracolonial conflicts, eventually leads to complete intraspecific worker-loss: a state where larvae have lost their developmental plasticity and develop as workers or queens depending only on whether they are the product of hybridization or not, respectively.

### 4.1 Model limitations

Of course, our analyses are based on several idealized assumptions. In particular, we assumed that the probability for larvae to develop as workers is under complete larval genetic control. Typically the developmental fate of female larvae also depends on various environmental factors created by adult colony members, such as food quality and quantity (Brian, 1956; Trible and Kronauer, 2017), or mechanical (Penick and Liebig, 2012) and chemical (Schwander et al., 2008; Penick et al., 2012) stimuli. The conclusions of our study apply as long as these environmental effects are held constant (or evolve more slowly than genetic caste determination). In this case, worker-loss would emerge via royal cheats that modify larval developmental reaction norm to environmental effects in such a way that their carriers are more likely to develop as queens (Hughes and Boomsma, 2008; Wolf et al., 2018). We also assumed that caste determination and hybridization evolve via rare mutations with weak additive effects at a single locus. These assumptions, which are typical to adaptive dynamics and kin selection approaches, have been extensively discussed elsewhere in a general context (Frank, 1998; Rousset, 2004; Geritz and Gyllenberg, 2005; Dercole and Rinaldi, 2008). In particular, all our results extend to the case where traits are determined by many genes, provided genetic variance in the population remains small (Charlesworth, 1990; Iwasa et al., 1991; Abrams et al., 1993). Where mutations have large additive or dominance effects, we expect more complex evolutionary dynamics, such as genetic polymorphism. These dynamics can nonetheless be straightforwardly investigated with the recurrence equations we derived (eq. A-4 in Appendix). However, our model cannot accommodate potential interaction effects among loci (i.e. epistasis). If a quantitative genetics analysis in *Temnothorax curvispinosus* supports that caste determination is influenced by additive effects in this species (Linksvayer, 2006), only epistatic effects were found in *Pogonomyrmex rugosus* (Schwander and Keller, 2008). It would therefore be relevant in the future to allow for a more a complex genetic basis of caste determination, including epistasis (in particular in the context of the evolution of unorthodox reproductive systems, see next section). Another important assumption we made is that hybrid larvae do not develop into fertile queens, for instance owing to hybrid incompatibilities (Trible and Kronauer, 2017). If fertile hybrid queens are produced regularly, evolution towards worker-loss like in our model is less likely to happen as hybrids no longer make a reliable source of workers. In ants at least, the idea that hybrid queens are rarely fertile is supported by the contrast between high frequency of interspecific mating on one hand, and weak genetic signals of interspecific gene flow on the other (Umphrey, 2006; Feldhaar et al., 2008). Finally, we focused in the main text on the case where colony productivity increases linearly with workers (i.e. the probability that a sexual survives until reproduction increases linearly with the number of workers). More realistically, the gain in productivity brought by one additional worker is likely to decrease with increasing workforce (Nonacs and Tobin, 1992; Reuter and Keller, 2001). Such diminishing returns tend to favor cheating because the indirect benefit of developing into a worker gets smaller as colony size increases (e.g. Reuter and Keller, 2001; Field and Toyoizumi, 2020). In line with this, we find that worker-loss evolves even more easily under diminishing compared to linear returns (Appendix B.2.3 and fig. S3).

### 4.2 An adaptive path to unorthodox reproductive systems?

Our result that sperm parasitism favours the emergence of worker-loss via the fixation of royal cheats may be relevant to unorthodox reproductive systems found in ants. Of particular interest is social hybridogenesis, whereby females produced through regular intra-lineage mating or thelytokous parthenogenesis develop into queens, while workers emerge from eggs fertilised by allospecific males (Helms Cahan et al., 2002; Helms Cahan and Keller, 2003; Anderson et al., 2006; Romiguier et al., 2017; Lacy et al., 2019; Kuhn et al., 2020). Such a striking system was first described just two decades ago in *Pogonomyrmex* harvester ants (Helms Cahan et al., 2002), and has since been found in several species spread across 4 genera (Helms Cahan and Keller, 2003; Romiguier et al., 2017; Lacy et al., 2019; Kuhn et al., 2020). If these observations suggest that social hybridogenesis has evolved independently multiple times, the evolutionary origins of this complex system remain poorly understood (Anderson et al., 2008; Schwander et al., 2010; Lavanchy and Schwander, 2019). One early suggestion is based on the hypothesis that worker development requires the combination of co-adapted alleles at key loci (i.e. requires epistatic interactions, Helms Cahan and Keller, 2003). According to this theory, worker-loss in hybridogenetic lineages would have originated in the random loss of such combinations during episodes of ancestral hybridization. Present hybridization would then have evolved to restore genetic combinations and epistatic interactions in F1-hybrids allowing for worker development.

Here, we have shown mathematically that social hybridogenesis could also result from additive genetic effects on caste development and queen-larvae conflicts within colonies. This theory, previously described verbally in Anderson et al. (2006) and Anderson et al. (2008), may help explain the multiple convergence towards social hybridogenesis because virtually every sexual eusocial species should experience queen-larvae conflicts over caste investment. Furthermore, because this path to social hybridogenesis does not depend on changes in the sympatric species whose sperm is parasitized, our model is relevant to both cases of asymmetrical (where the sympatric species produces workers through regular sex, as in *Solenopsis xyloni* for e.g.; Helms Cahan and Keller, 2003) and symmetrical social hybridogenesis (where the sympatric species also produces workers via hybridization, as in *Pogonomyrmex* harvester ants for e.g.; Anderson et al., 2006).

Our model may also be relevant to other unorthodox systems of reproduction such as those found in populations of *Wasmannia auropunctata* (Fournier et al., 2005), *Vollenhovia emeyri* (Ohkawara et al., 2006) or *Paratrechina longicornis* (Pearcy et al., 2011). As with some forms of social hybridogenesis, queens of these systems produce their reproductive daughters via female parthenogenesis and their workers via sex with genetically distant males. In contrast to social hybridogenesis, however, these males belong to a divergent all-male lineage maintained by male clonality. This is further accompanied with a complete absence of arrhenotokous males (i.e. queens never make hemiclonal haploid sons, as shown in *W. auropunctata*, Rey et al., 2013). When queens are able to produce daughters parthenogenetically in our model, evolution can lead to a state where worker-loss is coupled with a complete absence of intra-lineage mating (i.e. *η* = 1, fig. 5C-D). In this state, arrhenotokous males represent a genetic dead-end, laying the basis for their disappearance. To investigate these systems in more detail, it would be interesting to extend our model to consider the evolution of female parthenogenesis and male clonality.

Our formal approach is especially useful in a context where hybrid vigour in workers has been raised to explain the evolutionary origin of social hybridogenesis and other hybridization-dependent systems (Julian and Cahan, 2006; Umphrey, 2006; Anderson et al., 2008; Feldhaar et al., 2008; Schwander et al., 2010). According to this argument, selection favoured hybridization because hybrid workers are more efficient, more resilient, or better suited to exploit marginal habitats than regular workers. But in spite of much effort, empirical evidence supporting hybrid vigor in workers is still lacking (Robertson and Ross, 1990; Julian and Cahan, 2006; Feldhaar et al., 2008, but see James et al., 2002). Further challenging this view, we have shown here that hybrid vigor is not necessary to the evolution of hybridization-dependent reproductive systems. In fact, our results demonstrate that these systems can easily evolve even when hybridization is costly due to pre-and post-zygotic barriers (i.e. when *e* < 1 for e.g. because hybridization leads to an inefficient workforce due to hybrid incompatibilities in workers; or increased efforts in mate-finding and mating, Maroja et al., 2014; or the production of non-viable or infertile hybrid queens, Umphrey, 2006; Feldhaar et al., 2008). In contrast to previous suggestions (Anderson et al., 2008), our model thus indicates that hybridization-dependent reproductive systems can emerge among species that have already substantially diverged, and can be maintained even with further accumulation of hybrid incompatibilities.

More generally, our results suggest that natural selection can lead to an association between hybridization and caste determination. To date, such associations have been reported in only 18 distinct ant species or populations (Helms Cahan et al., 2002; Helms Cahan and Keller, 2003; Fournier et al., 2005; Anderson et al., 2006; Ohkawara et al., 2006; Pearcy et al., 2011; Romiguier et al., 2017; Lacy et al., 2019; Kuhn et al., 2020). But this rarity may be due – at least partly – to the difficulty with describing these systems (which in particular requires sampling and genotyping both queens and workers of the same populations, Helms Cahan et al., 2002). For instance, studies specifically testing for social hybridogenesis discovered 5 new cases of social hybridogenesis in *Cataglyphis* (out of 11 species tested, Kuhn et al., 2020) and 3 in *Messor* (out of 9, Romiguier et al., 2017). These considerations, together with our results, support the notion that currently known cases likely represent only a small fraction of extant eusocial systems relying on hybridization (Helms Cahan et al., 2002; Lavanchy and Schwander, 2019).

### 4.3 Factors promoting the evolution of intraspecific worker-loss

In addition to showing that hybrid vigor is not necessary for the emergence of intraspecific worker-loss, our model highlights several factors that can facilitate such evolution. The first of these is polyandry, which favors sperm parasitism and worker-loss by minimizing the risks associated with hybridization. Interestingly, even though polyandry is generally rare in social insects (Strassmann, 2001; Hughes et al., 2008), meaningful exceptions are found in *Pogonomyrmex* (Rheindt et al., 2004) and *Messor* (Norman et al., 2016) harvester ants, two taxa where social hybridogenesis has evolved multiple times (Anderson et al., 2006; Romiguier et al., 2017). While the number of males a queen mates with is fixed in our model, it is conceivable that this number also responds to hybridization, leading polyandry and hybridization to coevolve. Indeed since low levels of polyandry represent a risk for out-breeding queens, we can expect selection to favour queen behaviours that increase their number of mates. This would in turn allow for greater levels of hybridization, which would increase selection on polyandry and so on. We therefore expect that the coevolution between polyandry, hybridization, and caste determination further promotes the emergence of worker loss. For species that are fixed for strict (or close to) monandry, our model shows that worker-loss can evolve when queens have the ability to reproduce via thelytokous parthenogenesis as it allows interspecifically mated queens to nevertheless produce daughter queens. This supports the notion that thelytoky has been important for the convergent evolution of social hybridogenesis in the (mostly) monandrous *Cataglyphis* ants (Kuhn et al., 2020).

Although not considered in our study for simplicity, another factor that can minimize the risks associated with hybridization in monandrous species is polygyny, whereby related queens form multi-queen nests. Such social organization allows both intra-and interspecifically mated queens to be part of the same colony, which can then produce both queens and workers. Polygyny should there-fore further facilitate hybridization. While this may have played a role in the evolution of social hybridogenesis in the polygynous *Solenopsis* species with this reproductive system (Helms Cahan and Keller, 2003; Lacy et al., 2019), we do not expect polygyny to be critical for the evolution of worker-loss as such loss has been described in both monogynic and polygynic species of the same genus (e.g., *Messor barbarus* and cf. *structor*; Romiguier et al., 2017). Beyond these considerations, any trait (e.g. polyandry, polygyny or reproduction by workers) that influences kinship structure within colonies and thus modulates intra-colonial conflicts has the potential to play a role in the evolution of worker-loss. Studying the evolution of such traits and its feedback on hybridization and caste determination therefore represents an interesting avenue for future research.

More important for the evolution of worker-loss in our model is that queens hybridize often enough. This readily happens when the propensity of queens to mate with allo*vs*. con-specific males evolves (fig. 3). In this case, sperm parasitism, worker-loss and social hybridogenesis emerge even in species that initially do not hybridize. Such evolution of hybridization is especially likely to occur where queens are able to recognize differences among males and choose their mates accordingly. There is however currently little, if any, evidence for such direct mate or sperm choice in eusocial insects (Strassmann, 2001; Schwander et al., 2006; Umphrey, 2006; Feldhaar et al., 2008). Alternatively, queens may be able to modulate the degree of hybridization via more indirect mechanisms, such as mating flight synchronization (Kaspari et al., 2001). Under completely random mating, hybridization can reach sufficient levels for worker-loss to evolve in our model as long as allo-specific males are sufficiently abundant (fig. 2), for instance because phenology is shared with an ecologically dominant species (Klein et al., 2017). In intermediate situations where allo-specific males are available but scarce, the evolution of caste determination under random mating leads to a situation where queens produce both hybrid and non-hybrid workers (fig. 2A-B). Such a scenario may be relevant to species of ants where hybrid workers has been reported but where worker-loss has not evolved (e.g. in some North American *Solenopsis* or European *Temnothorax*, Feldhaar et al., 2008).

Whether it occurs randomly or not, hybridization requires pre-zygotic barriers to be sufficiently low. Various mechanisms, such as secondary contacts or high dispersal ability, are known to lower these barriers (Aguiar et al., 2009). In particular, it has been proposed that the typically low phenotypic variation among males of different ant species facilitates hybridization in this taxa (Feldhaar et al., 2008). With these considerations in mind, it is noteworthy that all known cases of social hybridogenesis have been found in ants that live in dry climates (Helms Cahan et al., 2002; Helms Cahan and Keller, 2003; Romiguier et al., 2017; Lacy et al., 2019; Kuhn et al., 2020), where the synchronicity of mating flights between species is highest due to shared dependence upon punctual climatic events (Hölldobler and Wilson, 1990; Feldhaar et al., 2008).

At a broader level, our results suggest that worker-loss can readily evolve when a source of workers that is impervious to royal cheats can be exploited by queens. Besides sperm parasitism, other forms of parasitism can provide such a source of workers and have been associated with worker-loss (Nonacs and Tobin, 1992). In inquiline ants such as *Teleutomyrmex schneideri* for instance, queens do not themselves produce workers but rather infiltrate the colony of a host and trick host workers into caring for their progeny (Hölldobler and Wilson, 1990; Buschinger, 2009). Like in our model, such social parasitism could be the endpoint of an arms race between queens and larvae of the same lineage, whereby increasingly caste-biasing cheats reduce colony workforce leading queens to increasingly rely on host workers.

### 4.4 Conclusions

Intra-colonial conflicts are inevitably part of the social lives of non-clonal organisms. Here we have shown that such genetic conflicts readily lead to an association between interspecific sperm parasitism and intraspecific worker-loss via the fixation of royal cheats. This association is especially relevant to the evolution of reproductive systems that like social hybridogenesis rely on hybridization. Beyond these unorthodox systems and sperm parasitism, the fixation of royal cheats and loss of intraspecific workers may be connected to other forms of antagonistic interspecific relationships such as social parasitism. More broadly, our model illustrates how the unique genetic conflicts that are inherent to eusocial life can lead to evolutionary arms races, with implications for elaborate re-productive systems and novel ecological interactions between species.

## Acknowledgements

We thank Nicolas Galtier for useful discussions, Miya Qiaowei Pan, Laurent Keller and Tanja Schwander for comments on an earlier version of our manuscript and the Swiss National Science Foundation (PCEFP3181243 to CM) for funding.

## Appendices

### A Methods

Here we describe our methods to investigate the evolutionary dynamics of: (1) the probability *ω* for a non-hybrid larvae to develop as a worker; and (2) the propensity *η* for queens to hybridize. These methods are organised as follows. First in section A.1, we present a population genetics model that describes the change in allele frequencies at a biallelic locus that determines the value of *ω* in larvae and of *η* in queens. Second (in section A.2.1), we obtain the invasion fitness of a mutant allele coding for deviant trait values in a population otherwise monomorphic for a resident allele. Then, we use this invasion fitness in section A.2 as a platform to infer the long-term adaptive dynamics of both traits (i.e. their gradual evolution under the input of rare mutations with weak phenotypic effects). Specifically, we derive the joint evolutionary equilibria of *ω* and *η* (i.e. singular values), as well as their properties (i.e. convergence stability and evolutionary stability, Dercole and Rinaldi, 2008 for textbook treatment). Finally in section A.3, we describe our individual-based simulations. A Mathematica notebook reproducing our analyses and figures is provided as a supplement here: https://zenodo.org/record/4434257.

#### A.1 Short term evolution: population genetics

##### A.1.1 Set-up

We consider a single locus with two alleles, *a* and *b*, that affect the expression of both *ω* and *η* in their carrier. Specifically, the probability for a larva with genotype *v* ∈ {*aa, ab, bb*} to develop as a worker is *ω*_*v*_, while each mate of a queen with genotype *v* ∈ {*aa, ab, bb*} is allospecific with a probability *η*_*v*_. To track the segregation of alleles *a* and *b* in the population, we let 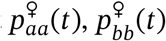, and 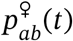 respectively denote the proportion of queens with genotype *aa, bb* and *ab* before mating at generation *t* (with 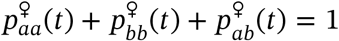). Similarly, 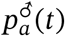 and 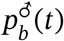 respectively denote the proportion of conspecific males with haploid genotype *a* and *b* before mating at generation *t* (with 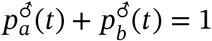).

##### A.1.2 Recurrence equations for the evolution of genotype frequencies

Our first goal is to develop recurrence equations for the frequencies of each genotype in males and females (i.e. express 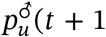 and 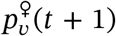 in terms of 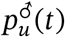 and 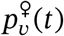 for *u* ∈ {*a, b*} and *v* ∈ {*aa, ab, bb*}). By definition, these frequencies can be written as

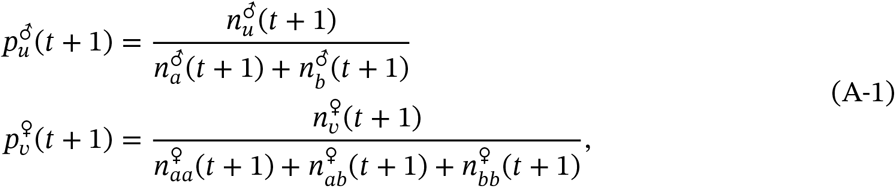

where 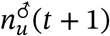 is the number of males of genotype *u* ∈ {*a, b*} at generation *t*+ 1, and 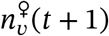 the number of queens of genotype *v* ∈ {*aa, ab, bb*} at generation *t* + 1 in the mating pool. Under our assumption that the probability for a sexual to reach the mating pool increases with the workforce of a colony (section 2 in main text), the numbers of males and females of each genotype can be expressed as:

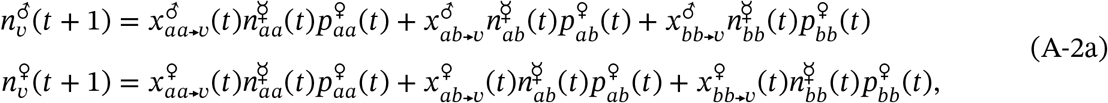

where 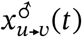 is the number of males with genotype *v* ∈ {*a, b*}, and 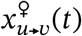 the number of queens with genotype *v* ∈ {*aa, ab, bb*}, produced by a colony founded by a queen of genotype *u* ∈ {*aa, ab, bb*} at generation *t*. Following Reuter and Keller (2001), we assume that these numbers are proportional to the energy invested into the production of sexuals. So instead of numbers, 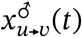 can be viewed as the investment into the production of males (of genotype *v* ∈ {*a, b*}) and 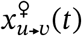 into the production of queens (of genotype *v* ∈ {*aa, ab, bb*}) by a colony whose queen has genotype *u* ∈ {*aa, ab, bb*}. Finally, 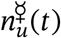 is the effective workforce of a colony whose queen has genotype *u* ∈ {*aa, ab, bb*} at generation *t*. This effective workforce is given by the sum of all types of workers present in a colony, including hybrids (with the latter weighted by their efficiency *e*), i.e.

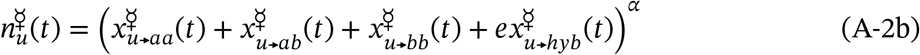

where 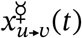 is the investment into the production of workers of genotype *v* ∈ {*aa, ab, bb, hyb*} (with *hyb* denoting hybrid genotype) made by a colony whose queen has genotype *u* ∈ {*aa, ab, bb*} at generation *t*. The parameter *α* > 0 determines the effect of the workforce on the probability for a sexual to reach the mating pool. When *α* = 1, investment in workers affects the survival of queens and males linearly (i.e. one extra unit of workforce always increases survival by the same amount). By contrast when *α* < 1, investment in workers show diminishing returns. Conversely when *α* > 1, investment in workers show increasing returns. For most of our analyses, we assume linear effects of the workforce (*α* = 1). We relax this assumption in section B.2.3.

We specify the investments into males, 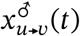, queens, 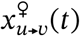, and workers, 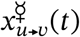, in terms of model parameters in Table S1. For the sake of completeness, we do so for a model that encompasses all the effects explored sequentially in the main text, i.e. we allow for both traits *ω* and *η* to coevolve; for a finite number *m* of mates for each queen; and for a fraction *c* of a queen’s brood to be produced via parthenogenesis. To read Table S1, note that the different investments made by a colony with a queen of type *u* ∈ {*aa, ab, bb*} (i.e. 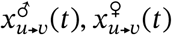, and 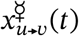) depend on the types of males she has mated with. To capture this, we let *M*_*u,v*_ be the random number of males of genotype *v* ∈ {*a, b, h*} (where *h* denotes allospecific type) that a queen of genotype *u* ∈ {*aa, ab, bb*} mates with. Assuming that each mate is independent from one another, these random variables follow a multinomial distribution with parameters,

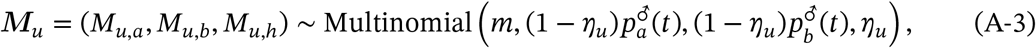

where *m* is the total number of mates; 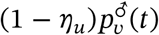 is the probability that in one mating event a queen of type *u* mates with a conspecific male of type *v* ∈ {*a, b*} (which requires that this queen does not hybridize, with probability (1 −*η*_*u*_), and encounters a male of type *v*, with probability given by its proportion, 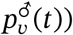; and *η*_*u*_ is the probability that in one mating event a queen of type *u* mates with an allospecific male.

To get to the recurrence equations tracking the proportion of males and queens of each genotype, we first substitute the entries of Table S1 into eq. (A-2) (with *α* = 1). Doing so we obtain polynomials for the numbers 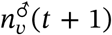 (for *v* ∈ {*a, b*}) and 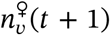 (for *v* ∈ {*aa, ab, bb*}) in terms of the random variables *M*_*u,a*_, *M*_*u,b*_, and *M*_*u,h*_ (with *u* ∈ {*aa, ab, bb*}). We marginalise (i.e. take the expectation of) these polynomials over the joint probability mass function of *M*_*u,a*_, *M*_*u,b*_, and *M*_*u,h*_ for each *u* ∈ {*aa, ab, bb*}, which is given by eq. (A-3). Finally, the so-obtained numbers of different types of individuals (eq. A-2) are substituted into eq. (A-1). From this operation and using the fact that 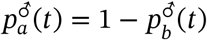 and 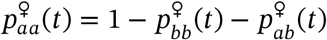, we obtain a recurrence equation,

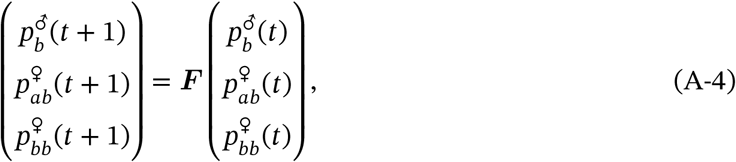

that is characterised by a mapping ***F*** : [0, 1]^3^ → [0, 1]^3^. This recurrence is too complicated to be presented here for the general case but can straightforwardly be iterated numerically to track allelic frequency changes for given parameter values (see Mathematica notebook for e.g.).

**Table S1:**
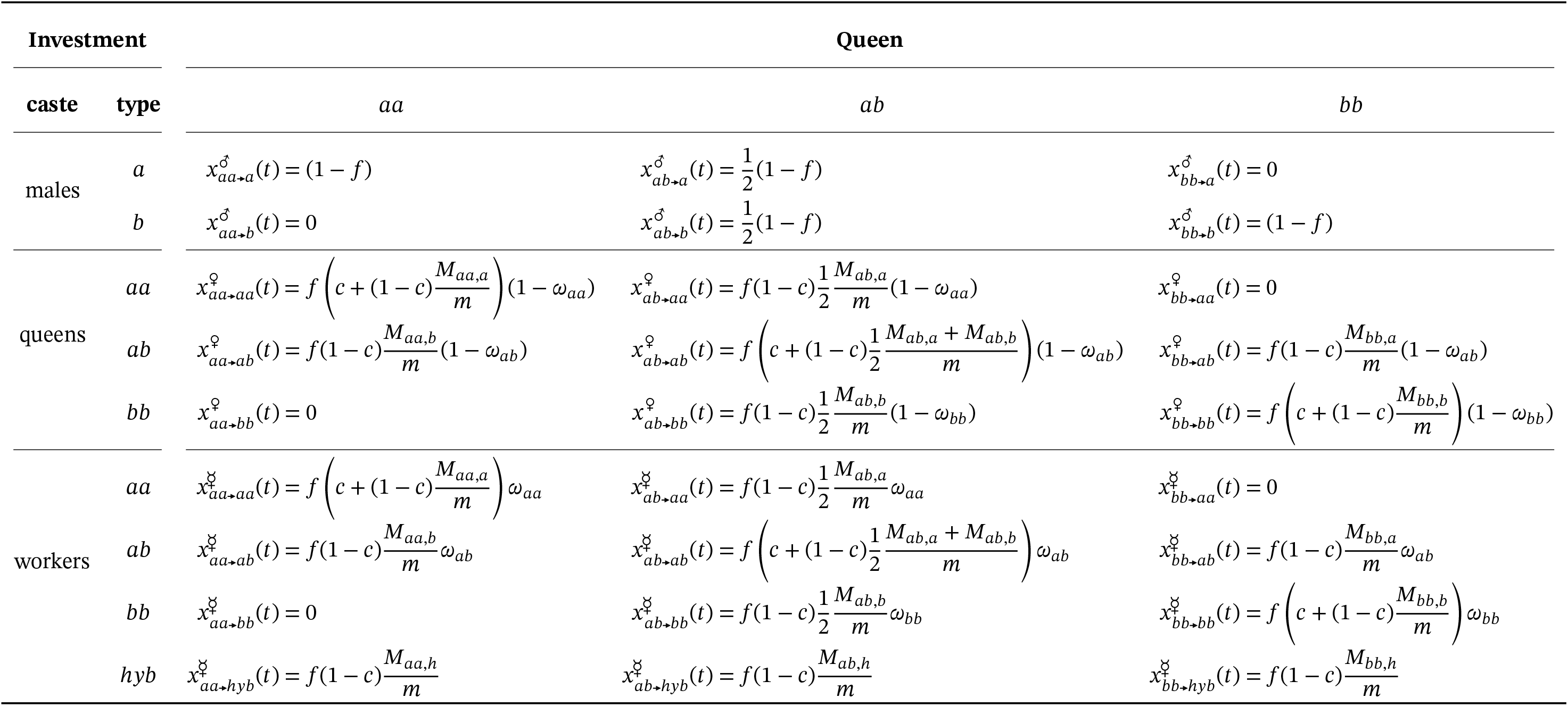
Colonial investment in males, queens and workers. Each entry in the table gives the investment into one type of individuals (given caste/genotype combination; rows), in a colony led by a queen with a given genotype (columns). Each expression depends only on models parameters and genotypic values for each trait. Genotypic values for hybridization probability in queens of genotype *u* (*η*_*u*_) do not appear explicitly but determine the distribution of the random variables *M*_*u,a*_, *M*_*u,b*_ and *M*_*u,h*_ (eq. A-3). To see how we constructed this table, consider for e.g. the investment 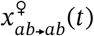 in queens of genotype *ab* in a colony led by a queen of genotype *ab* (fourth row, second column). First, queens can arise only from the fraction *f* of the brood that is diploid. Next, as the laying queen is of genotype *ab*, the fraction *c* of diploid eggs that are produced through parthenogenesis will also be of genotype *ab*. The fraction (1 − *c*) of diploid eggs that are fertilised through regular sex is *ab* with a probability that depends on the queen’s mates: (*M*_*ab,a*_ + *M*_*ab,b*_)/(2*m*) (assuming random chromosomal segregation and fertilisation, e.g. because the amount of sperm provided by each male is the same and well-mixed within a queen’s spermathecae). Finally, diploid *ab* eggs develop into queens with probability 1− *ω*_*ab*_. The other entries of the table are derived similarly.

#### A.2 Long-term evolution: adaptive dynamics

To gain more analytical insights, we use the recurrence eq. (A-4) to study the long term adaptive dynamics of both traits under the assumption that traits evolve via mutations that are rare and with weak additive phenotypic effects.

##### A.2.1 Invasion fitness of rare additive allele

An adaptive dynamics model is typically based on the invasion fitness of a mutant allele in a population that is otherwise fixed for a resident allele (i.e. the asymptotic growth rate of a mutant allele). To obtain this invasion fitness, we first introduce some notation. We denote the resident allele by a vector ***z*** = (*ω, η*) where *ω* is probability that a larva homozygote for the resident allele develops into a worker, and *η* is the probability that a mate of queen homozygote for the resident allele is allo-specific. Similarly, the mutant allele is described by a vector ***ζ*** = (*ω* + *δ*_*ω*_, *η* + *δ*_*η*_) whose first entry gives the probability that a larva homozygote for the mutant allele develops into a worker, and whose second entry is the probability that a mate of a queen homozygote for the mutant allele is allo-specific (*δ*_*ω*_ and *δ*_*η*_ thus denote the mutant effect on trait values). Assuming additive genetic effects on phenotypes, a heterozygote then expresses phenotype (*ω* + *δ*_*ω*_/2, *η* + *δ*_*η*_/2).

To use the recurrence equations developed in the previous section, we arbitrarily set allele *a* as the resident and *b* as the mutant. The allele specific trait values (appearing in Table S1 and eq. A-3) are then replaced by:

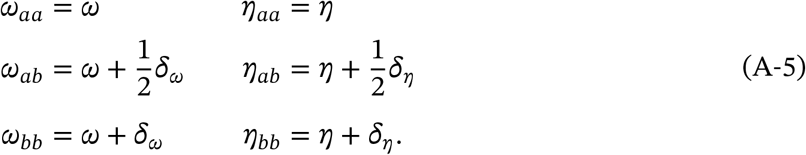

Next, we use the fact that the mutant is rare so that its frequency in the population is of the order of a small parameter denoted 0 < *ϵ* ≪ 1. As a rare allele can only be found in heterozygous form in a large panmictic population, the initial dynamics of a mutant allele *b* can be described through linear approximations of 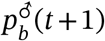 and 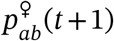 at a near-zero frequency of *b* (e.g. Brännström et al., 2013). In other words, eq. (A-4) can be linearised to

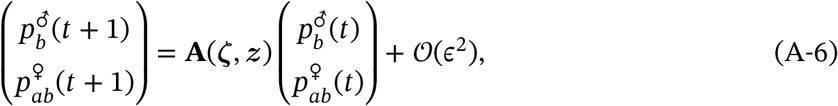

where **A**(***ζ, z***) is a 2×2 matrix that depends on mutant and resident phenotypes, ***ζ*** and ***z***, and *ϵ* is a small parameter of the order of the frequency of the mutant *b* in males and queens.

The invasion fitness of the mutant phenotype, which we write as *W*(***ζ, z***), is then given by the leading eigenvalue of **A**(***ζ, z***) (e.g. Caswell, 2000), i.e.

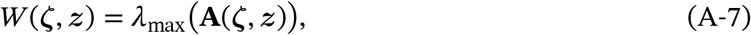

where *λ*_max_ (**M**) gives the leading eigenvalue of a matrix **M**. In a large population, *W*(***ζ, z***) tells the fate of the mutant allele. If *W*(***ζ, z***) ≤ 1, then the mutant allele is purged by selection and vanishes with probability one. Otherwise if *W*(***ζ, z***) > 1, the mutant has a non zero probability of invading the population (e.g. Brännström et al., 2013).

##### A.2.2 Directional selection

When mutations are rare with weak phenotypic effects, the population first evolves under directional selection whereby an advantageous mutation fixes before a new mutation arises so that the population “jumps” from one monomorphic state to another (Dercole and Rinaldi, 2008). To study these dynamics, we use the selection gradient, ***s***(***z***), which is a vector pointing in the direction favoured by selection at every point ***z*** ∈ [0, 1]× [0, 1] of the phenotypic space (i.e., the space of all possible phenotypic combinations with *ω* and *η* both between 0 and 1 as they are both probabilities) . This vector is given by the marginal effect of each trait on invasion fitness, i.e.

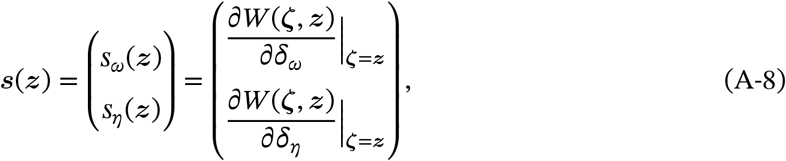

where *s*_*ω*_(***z***) and *s*_*η*_(***z***) give the direction of selection on *ω* and *η* respectively.

###### Singular strategies

A singular strategy, ***z***^∗^ = (*ω*^∗^, *η*^∗^), is such that all selection gradients are equal to zero,

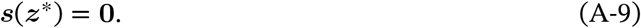

A singular strategy therefore represents a potential equilibrium of adaptive dynamics (Brännström et al., 2013).

###### Jacobian matrix and convergence stability

Whether the population evolves towards or away from a singular strategy ***z***^∗^ depends on the Jacobian matrix,

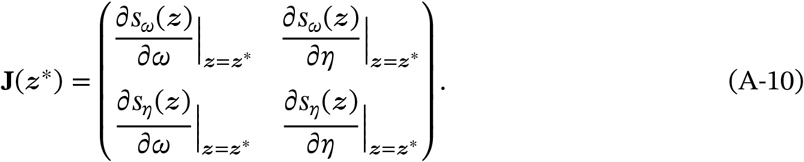

Specifically, one necessary condition for a singular strategy to be an evolutionary attractor is that the greatest real part of the eigenvalues of **J**(***z***^∗^) is negative (Leimar, 2009). Such a singular strategy ***z***^∗^ is said to be convergence stable. Otherwise, the population will be repelled away from ***z***^∗^. Even if ***z***^∗^ is convergence stable, it is possible for the population to evolve away from ***z***^∗^ when both evolving traits are genetically correlated (Leimar, 2009). A sufficient condition for a singular strategy to be an attractor is that the symmetric part of the Jacobian matrix, (**J**(***z***^∗^)+**J**(***z***^∗^)^T^)/2, is negative-definite, in which case ***z***^∗^ is said to be *strongly* convergence stable (Leimar, 2009). When this is true, the population evolves towards ***z***^∗^, whatever the genetic correlations between both traits (i.e. independently from the statistical distribution of mutational effects on both traits).

##### A.2.3 Stabilising/disruptive selection

Once the population is at an equilibrium for directional selection (i.e. a convergence stable phenotype), it either remains monomorphic under stabilising selection (when the equilibrium is evolutionary stable or uninvadable, Parker and Maynard Smith, 1990) or becomes polymorphic due to disruptive selection (when the equilibrium is not evolutionary stable or invadable, Geritz et al., 1998). When two traits are coevolving, this depends on the Hessian matrix (Phillips and Arnold, 1989; Leimar, 2009; Geritz et al., 2016),

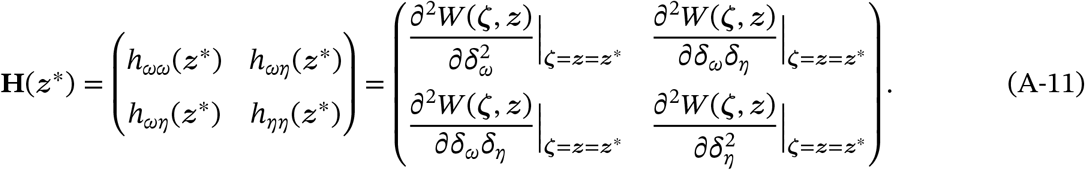

An equilibrium ***z***^∗^ is uninvadable if **H**(***z***^∗^) is negative-definite. Otherwise, selection may be disruptive and the population may experience evolutionary branching, whereby it splits among two diverging morphs (Geritz et al., 1998; Leimar, 2009; Geritz et al., 2016).

#### A.3 Individual-based simulations

To complement our mathematical analysis, we also performed individual based simulations (an R script implementing these is provided as a supplement here: https://zenodo.org/record/4434257). These simulations track a population of *N*_*q*_ = 10000 diploid queens (with *f* = 0.5, see figure legends for other parameters). Each queen is characterized by its genotype: a pair of haplotypes, each of which is given by the values of *ω* and *η* they code for (so four genotypic values in total). Simulations are initialized by setting both haplotypes of all *N*_*q*_ queens to the same arbitrary values (i.e. we start with a monomorphic population). Each generation of a simulation consists of the following steps:

1. **Mating**. First, queens mate. To model this process, we first compute the propensity *η*_*i*_ of each queen *i* ∈ {1, 2,…, *N*_*q*_} to hybridize as the mean of the two relevant alleles it is carrying. Then, each queen *i* is mated with a number *m*_*i*_ of conspecific haploid males. This number *m*_*i*_ is drawn from a binomial distribution with *m* trials and success probability (1−*η*_*i*_) (in line with eq. A-3). At the first generation, all males carry the same genetic values for *ω* and *η* as queens (i.e. the initial trait values). In subsequent generations, males are sampled (with replacement) as single haplotypes from the 2*i* haplotypes present in the laying queens of the previous generation. Following eq. (A-2a), the probability to sample a given haplotype is weighted by the investment in workers within its colony of origin (as the investment in workers increases the probability for males to reach the mating pool).
2. **Colony development**. Each queen *i* settles to form a colony. We characterise each colony in two steps. First, a list is constructed that contains the 2*m*_*i*_ non-hybrid diploid female genotypes produced within each colony (i.e. the combinations of the alleles of a queen and of its conspecific mates). If thelytokous parthenogenesis is included (*c* > 0), the genotype of the queen itself is added to this list. Second, the investment in workers within each colony is calculated following equations in Table S1 and eq. (A-2b). These calculations use the genetic value expressed by each of the 2*m*_*i*_ +1 non-hybrid genotype within the female progeny (characterised in the first step), as well as the proportion of the brood produced sexually and asexually (the parameter *c*), the proportion of conspecific and allospecific males the queen has mated with (i.e. *m*_*i*_/*m* and 1− *m*_*i*_/*m*), and the efficiency of hybrid workers (the parameter *e*).
3. **Next-generation queens**. To generate the next generation of queens, *N*_*q*_ new diploid female genotypes are sampled (with replacement) from all non-hybrid genotypes produced within each colony. Following Table S1, the probability to sample a given genotype is weighted by its own genetic value of (1 − *ω*) and by the investment in workers within its colony of origin (as the investment in workers increases the probability for queens to reach the mating pool). Finally, each genotypic value independently mutates with probability 10^−2^. Mutation effects are drawn independently from a normal distribution with mean 0 and standard deviation 10^−2^. Mutated genetic values are capped between 0 and 1 to ensure that traits remain within their domain of definition.

### B Analyses

Here, we present the derivations of our results summarised in the main text. These derivations are organised in the same order as they appear in the main text. As a supplement, we also provide a Mathematica (Wolfram Research, 2020) notebook that allows to follow our analyses.

#### B.1 Baseline model

We first explore the baseline case where females mate with a large (effectively infinite) number of mates and there is no parthenogenesis (i.e. when *m* → ∞ and *c* = 0).

##### B.1.1 Independent evolution of genetic caste determination

As presented in the main text, we initially assume that hybridization *η* is fixed and only caste determination *ω* evolves. Using eq. (A-8) with *m* → ∞ and *c* = 0, we find that the selection gradient on genetic caste determination is,

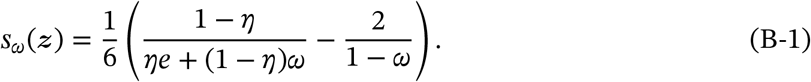

Accordingly, there is a unique singular strategy *ω*^∗^ for caste determination when hybridization *η* is fixed (i.e. *ω*^∗^ such that *s*_*ω*_((*ω*^∗^, *η*)) = 0),

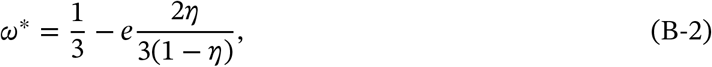

which is eq. 1 of the main text.

It is straightforward to show that with hybridization fixed, the singular strategy (eq. B-2) is convergence stable (plugging eq. B-2 into the Jacobian, that is eq. A-10, for a single trait with *m* → ∞ and *c* = 0),

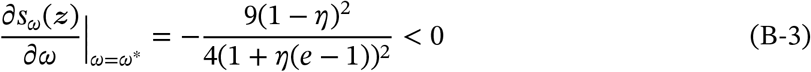

as well as uninvadable (plugging eq. B-2 into the Hessian, that is eq. A-11, for a single trait with *m* → ∞ and *c* = 0),

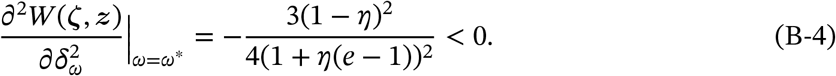

Therefore, when hybridization is fixed, our analyses show that genetic caste determination will gradually evolve to the singular value eq. (B-2) and remain monomorphic for this value (which is what we observe when we simulate this scenario, fig. 2A).

##### B.1.2 Coevolution of genetic caste determination and hybridization

###### An unstable singularity

When both caste determination *ω* and hybridization *η* evolve, their trajectories under directional selection are given by the selection gradient vector,

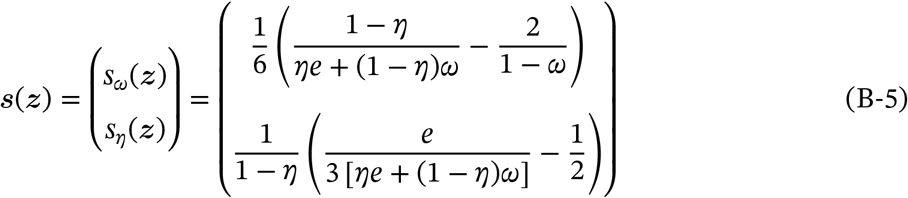

(from eq. A-8 with *m* → ∞ and *c* = 0). Solving the above for ***z***^∗^ = (*ω*^∗^, *η*^∗^) such that ***s***(***z***^∗^) = (0, 0) yields a single singular strategy in two dimensional trait space,

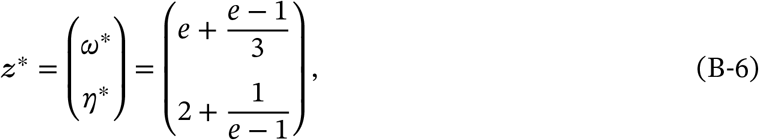

which is plotted in fig. 3A against *e*. However, when we look at the Jacobian matrix of the system eq. (B-5) at this singular value (i.e. substitute eqs. B-5 and B-6 into eq. A-10),

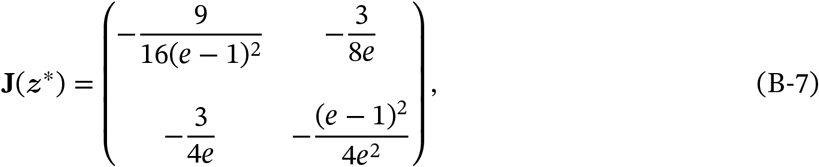

we see that this matrix has a negative determinant,

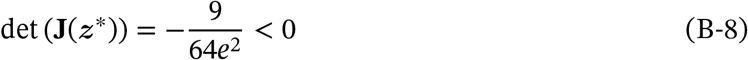

so its eigenvalues cannot both be negative (since the product of the eigenvalues of a matrix is equal to its determinant). Hence the singular value ***z***^∗^ eq. (B-6) is not convergence stable, but rather an evolutionary repellor.

Our result that evolutionary trajectories will be repelled away from the singular value eq. (B-6) tells us that adaptive dynamics will eventually get to the boundary of the trait space. This trait space consists of the square [0, 1] × [0, 1] (as both traits must be between zero and one). Two edges of this square (when *ω* = 1 or *η* = 1) cannot be accessed by evolutionary dynamics as either of these trait values lead to zero fitness (as a population monomorphic for *ω* = 1 or *η* = 1 produces no queen in our baseline model). We can therefore focus on dynamics along the edges *η* = 0 or *ω* = 0 of the trait space, which respectively correspond to the case of hybridization avoidance and worker-loss.

###### Convergence to hybridization avoidance

Evolutionary dynamics will settle somewhere on the edge where hybridization is absent in the population (*η* = 0) only if: (1) selection on hybridization maintains it at zero (i.e. *s*_*η*_(***z***) ≤ 0 when *η* = 0); and (2) selection on caste determination settles for an equilibrium *ω*^∗^ (i.e. *s*_*ω*_(***z***) = 0 for some *ω*^∗^ when *η* = 0). From eq. (B-5), these two conditions are true when *e* ≤ 1/2 and the equilibrium for caste determination is simply *ω*^∗^ = 1/3 (in line with eq. B-2). As established in eq. (B-3), this equilibrium is convergence stable and evolutionary stable when *η* is fixed.

###### Convergence to worker-loss

Similarly, for adaptive dynamics to converge somewhere on the edge where workers are no longer produced from regular sex (*ω* = 0), these two conditions are necessary: (1) selection on caste determination maintains *ω* = 0 (i.e. *s*_*ω*_(***z***) ≤ 0 when *ω* = 0); and (2) selection on hybridization favours an equilibrium *η*^∗^ (i.e. *s*_*η*_(***z***) = 0 for some *η*^∗^ when *ω* = 0). Substituting eq. (B-5) into these conditions, they reduce to *e* ≥ 1/4 and *η*^∗^ = 2/3. In addition, we see from eq. (B-5) that when *ω* = 0,

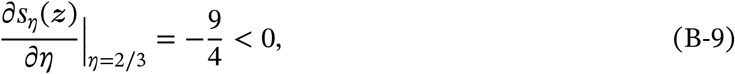

and we further find that

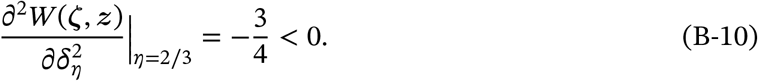

This tells us that the population will converge towards and remain monomorphic for *η*^∗^ = 2/3 when *ω* = 0 is fixed.

###### Three phase portraits

Put together, the above observations allow us to deduce that depending on the parameter *e*, there are three possible types of phase portraits for the adaptive dynamics of both traits (fig. 3B-D). When *e* ≤ 1/4, the singular value eq. (B-6) is outside of the trait space (or on its boundary when *e* = 1/4) and the point (*ω* = 1/3; *η* = 0) is an evolutionary stable attractor, meaning that the population will converge towards hybridization avoidance (fig. 3B). When *e* ≥ 1/2, the singular value eq. (B-6) is also outside of the trait space (or on its boundary when *e* = 1/2) and the point (*ω* = 0; *η* = 2/3) is an evolutionary stable attractor, meaning that the population will converge towards worker-loss (fig. 3D). Finally when 1/4 < *e* < 1/2, the singular value eq. (B-6) is a repellor that lies within the trait space (i.e. 0 < *ω*^∗^ < 1 and 0 < *η*^∗^ < 1) and both points (*ω* = 1/3; *η* = 0) and (*ω* = 0; *η* = 2/3) are evolutionary stable attractors. In this case evolutionary dynamics will depend on initial values (fig. 3C).

##### B.1.3 Decomposition of directional selection in terms of inclusive fitness effects

###### The kin selection approach

In this section, we use the so-called “kin selection” or “inclusive fitness” approach to obtain the selection gradient eq. (B-5) (Taylor and Frank, 1996). This approach, which is based on invasion analyses of alleles in class-structured populations, gives the same quantitative result about directional selection than other common methods in theoretical evolutionary biology such as adaptive dynamics, population or quantitative genetics (assuming genetic variance for traits is small, e.g. Taylor and Frank, 1996; Rousset, 2004; Lehmann et al., 2016). But one particular advantage of a kin selection approach is that it immediately decomposes directional selection on mutant alleles into the sum of: (1) their direct fitness effects on the reproductive success of the individuals that express them; and (2) of their indirect fitness effects on other related individuals that can also transmit them. This decomposition allows to delineate the various forces at play in the evolution of social behaviours (Hamilton, 1964). Here, we use it to better understand the evolution towards worker-loss (and obtain fig. 3E-F).

We follow Taylor and Frank (1996)’s general method. Consider a population with mean trait values *ω* and *η*. In this population, consider a focal colony that is home to a mutant allele that codes for deviant trait values *η*_•_ in queens and *ω*_•_ in larvae that carry this allele. Let *ω*_0_ denote the mean trait value expressed by all larvae within this focal colony. Using this notation, the expected number of successful (i.e. that mate) males that are produced by the focal colony and that carry the mutant allele is given by,

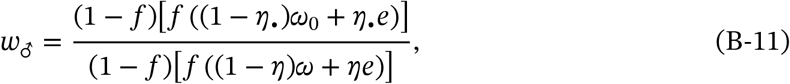

where the numerator and denominator are the total number of males produced by the focal and a random colony, respectively. For the focal colony (the numerator), (1 − *f*) is the probability that an egg is haploid (i.e. male) while the term in square brackets is the colony’s investment in workers (which in our model is also the probability that a sexual reaches maturity). The denominator follows the same logic for an average colony in the population.

Similarly, the expected number of successful queens that are produced by the focal colony that carry the mutant allele is,

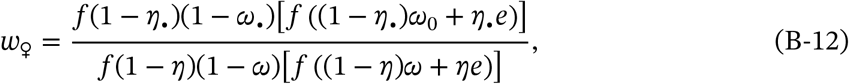

where *f*(1 −*η*_•_)(1 −*ω*_•_) is the number of queens produced in the focal colony and the term in square brackets is the probability that a queen survives till mating (i.e. the colony’s investment in workers).

###### Fitness effects within a mutant colony

With the above notation, the selection gradient vector can then be computed as,

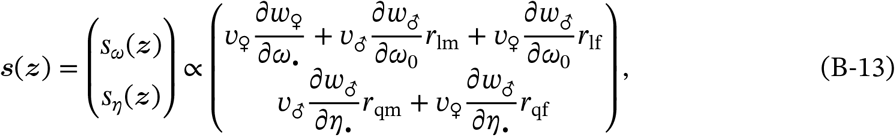

where all derivatives are evaluated at *ω*_•_ = *ω*_0_ = *ω*and *η*_•_ = *η*_0_ = *η*; *r*_lm_ is the relatedness of a female larva to a brother; *r*_lf_ is the relatedness of a female larva to a sister; *r*_qm_ is the relatedness of a queen to its sons; *r*_qf_ is the relatedness of a queen to its daughters; *v*_♂_ is the reproductive value of males and *v*_♀_ is the reproductive value of queens (all these relatedness coefficients and reproductive values are for a monomorphic population, Taylor and Frank, 1996; Rousset and Ronce, 2004; Lehmann et al., 2016). Plugging eqs. (B-11) and (B-12) into eq. (B-13) with relatedness coefficients and reproductive values corresponding to a haplodiploid system with infinite matings (i.e. *r*_lm_ = 1/2, *r*_lf_ = 1/4, *r*_qm_ = 1, *r*_qf_ = 1/2, *v*_♂_ = 1/2, *v*_♀_ = 1), we obtain expressions equivalent to eq. (B-5). But in contrast to eq. (B-5), the selection gradients in eq. (B-13) are expressed as a sum of fitness effects of a mutant allele via a given category of individual. More specifically, the gradient *s*_*ω*_(***z***) in eq. (B-13) is decomposed as the fitness effects of an allele coding for a mutant value of *ω* in larvae: on the larvae that express this allele (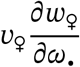, yellow line in fig. 3E), on their brothers 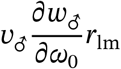, blue line in fig. 3E), and on their sisters (i.e. queens, 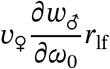, red line in fig. 3E) that can also transmit the allele. Similarly, the gradient *s*_*η*_(***z***) in eq. (B-13) is composed of the fitness effects of an allele coding for a mutant value of *η* in queens: via their sons 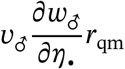, blue line in fig. 3F) and daughters (i.e. queens, 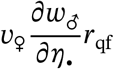, red line in fig. 3F). To construct panels E and F of fig. 3, we evaluated these five terms outlined above at every step of an evolutionary trajectory from the baseline equilibrium in absence of hybridization (*ω* = 1/3, *η* = 0) to complete worker-loss (*ω* = 0, *η* = 2/3). The evolutionary trajectory was obtained by iteration, starting from the baseline equilibrium and taking steps of size 0.001 (in units of trait space) in the direction of the selection gradient (eq. B-5).

##### B.1.4 Correspondence with Reuter and Keller (2001)

Here we connect our results to those of Reuter and Keller (2001), who used a kin selection approach to study the evolution of caste determination when under full queen, full larval, or mixed control (in the absence of hybridization). Our model corresponds to the case of full larval control (eq. 3 of Reuter and Keller, 2001). Our selection gradient *s*_*ω*_(***z***), shown in eq. (B-1) with *η* = 0, reduces to eq. 3 of Reuter and Keller (2001) when we assume linear effects of investment in workers on colony productivity. More specifically, if we set their term Δ_*c*_ = *δs*/(*δw*)× 1/*f* (their notation in their eq. 3, where Δ_*c*_ corresponds to the gain in sexual production brought by one additional worker) to

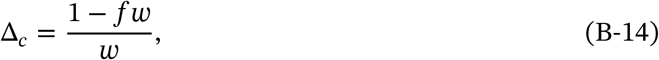

and assume that the population is monogynous and highly polyandrous with balanced sex-ratio (i.e. in their notation, *f* = 1/2; *g*_*f*_ = 1/4; *g*_*m*_ = 1/2; *v*_*f*_ = 2; *v*_*m*_ = 1), then we find that eq. 3 of Reuter and Keller (2001) is proportional to our selection gradient *s*_*ω*_(***z***) (eq. B-1) with *η* = 0. In line with this, both yield the convergence stable equilibrium *w*^∗^ = 1/3.

#### B.2 Extensions

We now consider several extensions to our baseline model.

##### B.2.1 Effect of finite matings

First, we relax our assumption that queens mate with an infinite number of mates (i.e. *m* < ∞).

###### Selection gradient

Working from eq. (A-8) with *c* = 0, we find that the selection gradient vector on caste determination *ω* and hybridization *η* under finite matings reads as,

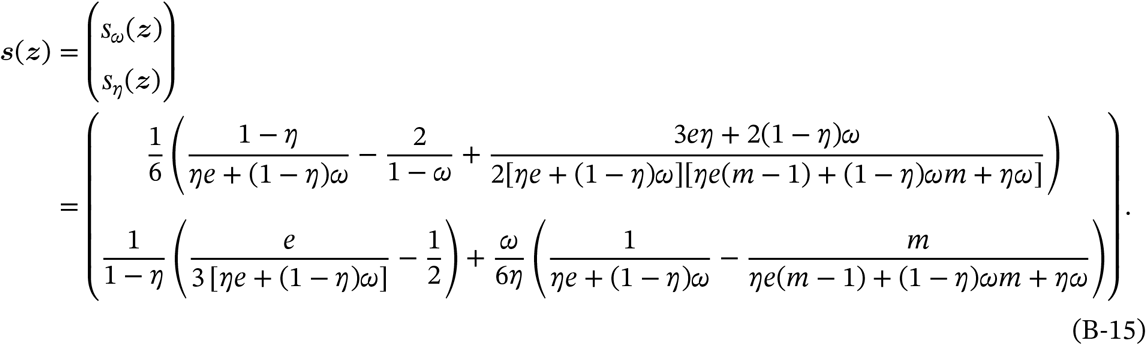

These gradients are complicated but we can extract relevant information by starting our analysis on the two boundaries of the trait space along which evolutionary dynamics may end up (*ω* = 0 or *η* = 0). Using eq. (B-15), we ask first when is worker-loss (*ω* = 0) stable? And second when is hybridization avoidance (*η* = 0) stable?

###### Stability of worker-loss

Worker-loss is stable only if: (1) selection maintains *ω* at zero (i.e. *s*_*ω*_(***z***) ≤ 0 when *ω* = 0); and (2) selection on hybridization settles for an equilibrium *η*^∗^ (i.e. *s*_*η*_(***z***) = 0 for some *η*^∗^ when *ω* = 0). From eq. (B-15), these two conditions reduce to

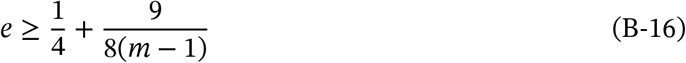

(region above dashed line in fig. 4A) and

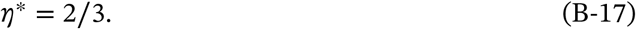

Note that condition (B-16) becomes impossible as *m* → 1. This indicates that worker-loss cannot evolve under monandry in this model. For *m* > 1, it is straightforward to show that when condition (B-16) is true, the strategy *η* = 2/3 is both convergence stable and evolutionary stable when *ω* = 0 (eqs. B-9 and B-10 for e.g. of the type of argument used).

###### Stability of hybridization avoidance

Conversely, hybridization avoidance is stable only if: (1) selection on hybridization maintains *η* at zero (i.e. *s*_*η*_(***z***) ≤ 0 when *η* = 0); and (2) selection on caste determination in absence of hybridization settles for an equilibrium *ω*^∗^ (i.e. *s*_*ω*_(***z***) = 0 for some *ω*^∗^ when *η* = 0). From eq. (B-15), these two conditions reduce to

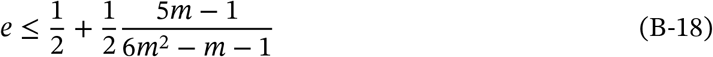

(region below plain line in fig. 4A) and

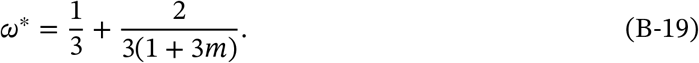

Again, it is straightforward to show that when condition (B-18) holds, the strategy given by eq. (B-19) is both convergence stable and evolutionary stable when *η* = 0 (eqs. B-3 and B-4 for e.g. of argument).

Together, conditions (B-16) and (B-18) split the parameter space into 4 areas where both, none, or only one of the conditions are met (fig. 4A). Where condition (B-18) is met but (B-16) is not (grey region of fig. 4A), hybridization cannot evolve when rare and worker-loss cannot be maintained. We therefore focus on the three remaining cases where worker loss can emerge. Doing so, we find that there are four possible types of evolutionary dynamics.

###### Type 1: Evolution towards worker-loss

Where condition (B-16) is met but (B-18) is not (dark green region of fig. 4A), selection favours the emergence of hybridization and maintenance of worker-loss. In addition, it can be shown that under these conditions, there exists no singular strategy within the trait space (i.e., there exists no ***z***^∗^ = (*ω*^∗^, *η*^∗^) such that 0 < *ω*^∗^, *η*^∗^ < 1 and ***s***(***z***^∗^) = (0, 0), e.g. using the function Reduce[] in Mathematica, see notebook). This means that the phase portrait of evolutionary dynamics is qualitatively the same as in fig. 3D: worker-loss always evolves.

###### Type 2: Evolution towards worker-loss or hybridization avoidance depending on initial conditions

Where conditions (B-16) and (B-18) are met simultaneously, both worker-loss and hybridization avoidance are stable so either strategy is maintained when common (when *m* ≥ 5, light green region of fig. 4A). Under these conditions, we find that there exists a singular strategy within the trait space (top row, columns *m* = 5 and *m* = 6 in fig. S1 for numerical values, see Mathematica notebook for analytical expression). When we compute numerically the leading eigenvalue of the system’s Jacobian matrix, we find that it is positive (fig. S1, second row, columns *m* = 5 and *m* = 6, dashed line), revealing that the singularity is an evolutionary repellor. Therefore the phase portrait of evolutionary dynamics is qualitatively the same as in fig. 3C: depending on initial conditions, evolutionary dynamics will lead to worker-loss or hybridization avoidance.

###### Type 3: Convergence stable and uninvadable intermediate strategy

Where neither condition (B-16) nor (B-18) are met, neither worker-loss nor hybridization avoidance are stable (when *m* ≤ 4, blue region of fig. 4A). In this case, a singular strategy within the trait space also exists (0 < *ω*^∗^, *η*^∗^ < 1; fig. S1, top row, columns *m* ∈ {1, 2, 3, 4} for numerical values; Mathematica notebook for analytical expression). But now, this intermediate strategy is (strongly) convergence stable as indicated by a negative leading eigenvalue of both the Jacobian matrix and its symmetric part (fig. S1, second row, columns *m* ∈ {1, 2, 3, 4}, dashed and dotted lines). When *m* ∈ {2, 3, 4}, this intermediate strategy is also uninvadable as shown by a negative leading eigenvalue of the Hessian matrix (fig. S1, second row, columns *m* ∈ {2, 3, 4}, full line). Thus, when the number of mates is between two and four (*m* ∈ {2, 3, 4}) and neither conditions (B-16) and (B-18) are met, the population converges and remains monomorphic for an intermediate strategy 0 < *ω*^∗^, *η*^∗^ < 1.

**Figure S1:**
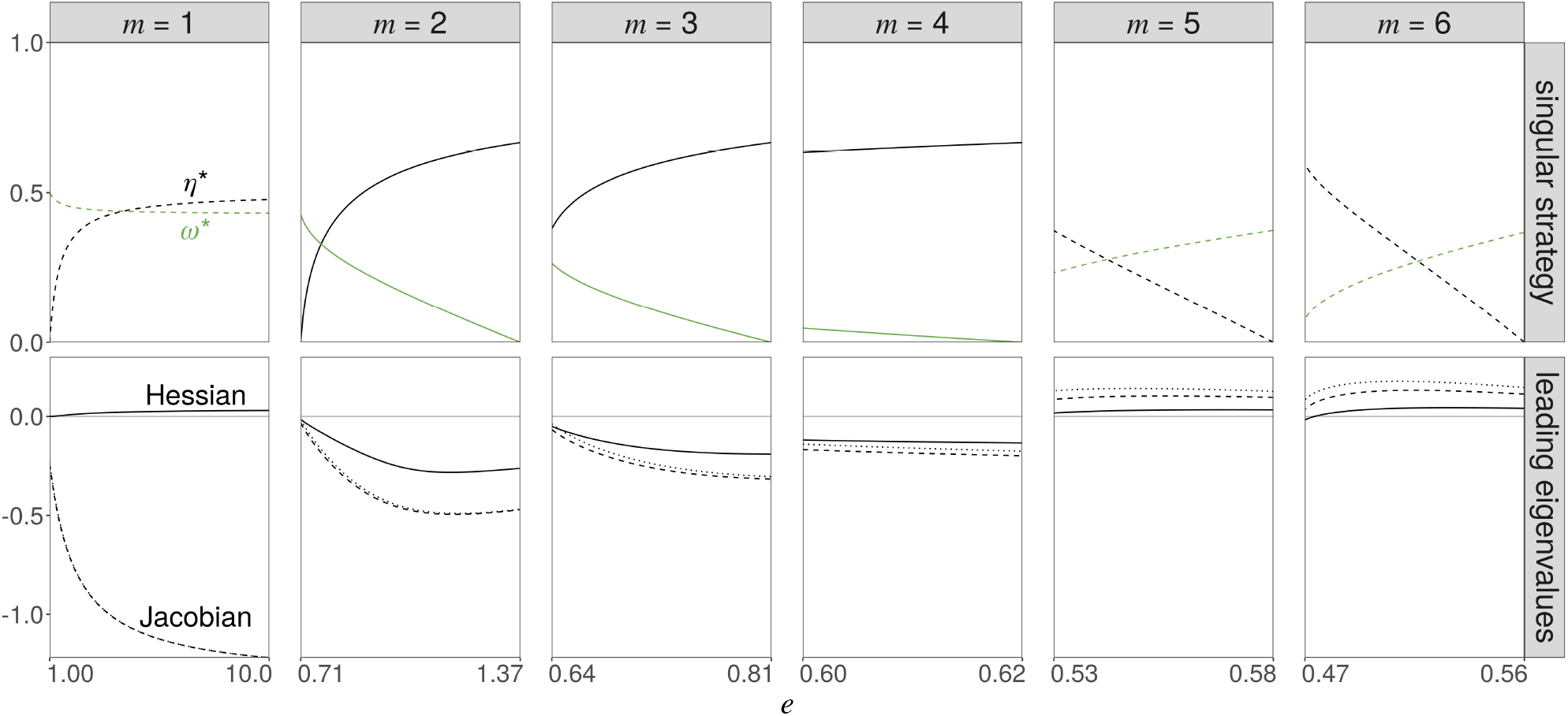
Properties of the internal singular strategy under monoandry and low polyandry. Each column describes the unique internal singular strategy for a specific value of *m*. **Top row**: value of the singular strategy (*ω*^∗^ in green, *η*^∗^ in black) within the range of *e* for which an internal strategy exists (range given by eqs. B-16 and B-18; Mathematica notebook for value of singular strategy). **Bottom row**: leading eigenvalues of the Jacobian (dashed line; for convergence stability), symmetric part of the Jacobian (dotted line; for strong convergence stability) and Hessian (full line; for evolutionary stability) matrices at the singular strategy (Mathematica notebook for calculations).

###### Type 4: Emergence of polymorphism under monandry

When neither condition (B-16) nor (B-18) are met and *m* = 1, the convergence stable intermediate strategy is invadable (i.e., the Hessian has a positive leading eigenvalue; fig. S1, second row, column *m* = 1, dashed line). This means that once the population has converged to this intermediate strategy, it experiences frequency-dependent disruptive selection leading to polymorphism (Geritz et al., 1998; Geritz and Gyllenberg, 2005; Geritz et al., 2016). Inspection of the entries of the Hessian matrix reveals that

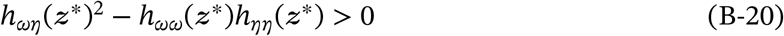

(fig. S2A, black line) and that *h*_*ωω*_(***z***^∗^) ≤ 0 and *h*_*ηη*_(***z***^∗^) ≤ 0 (fig. S2A, green and grey lines). This says that disruptive selection in our model is due to correlational selection between caste determination and hybridization (i.e. the selection that associates caste determination and hybridization, Phillips and Arnold, 1989) and only occurs because both traits are coevolving (i.e. if either trait evolves while the other is fixed, the population remains monomorphic, e.g. Mullon et al., 2018). We also find that

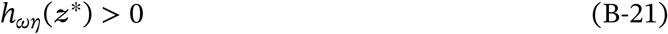

(fig. S2A, blue line), which tells us that correlational selection is positive (i.e. selection favours a positive correlation between caste determination and hybridization within individuals, Phillips and Arnold, 1989). This is confirmed by individual based simulations, in which we observe the emergence of a polymorphism characterised by a positive correlation between *ω* and *η* within haplotypes (fig. 4C and fig. S2B-D).

##### B.2.2 Effect of thelytokous parthenogenesis

When we allow for a fraction *c* of a queens brood to be produce parthenogenetically, the selection gradient (obtained from eq. A-8) is too complicated to be displayed or for singular strategies to be found analytically. We therefore go through an invasion analysis similar to above (Appendix B.1.2 and B.2.1) and again ask: (1) under which conditions and values of *ω* is hybridization avoidance (*η* = 0) stable? and (2) under which conditions and values of *η* is worker-loss (*ω* = 0) stable?

###### Stability of hybridization avoidance

Hybridization avoidance is stable if selection on caste determination settles for an equilibrium *ω*^∗^ in the absence of hybridization (i.e. *s*_*ω*_(***z***) = 0 for some *ω*^∗^

when *η* = 0), and if selection on hybridization at this equilibrium maintains *η* at zero (i.e. *s*_*η*_(***z***) ≤ 0 when *η* = 0 and *ω* = *ω*^∗^). These two conditions respectively reduce to,

**Figure S2:**
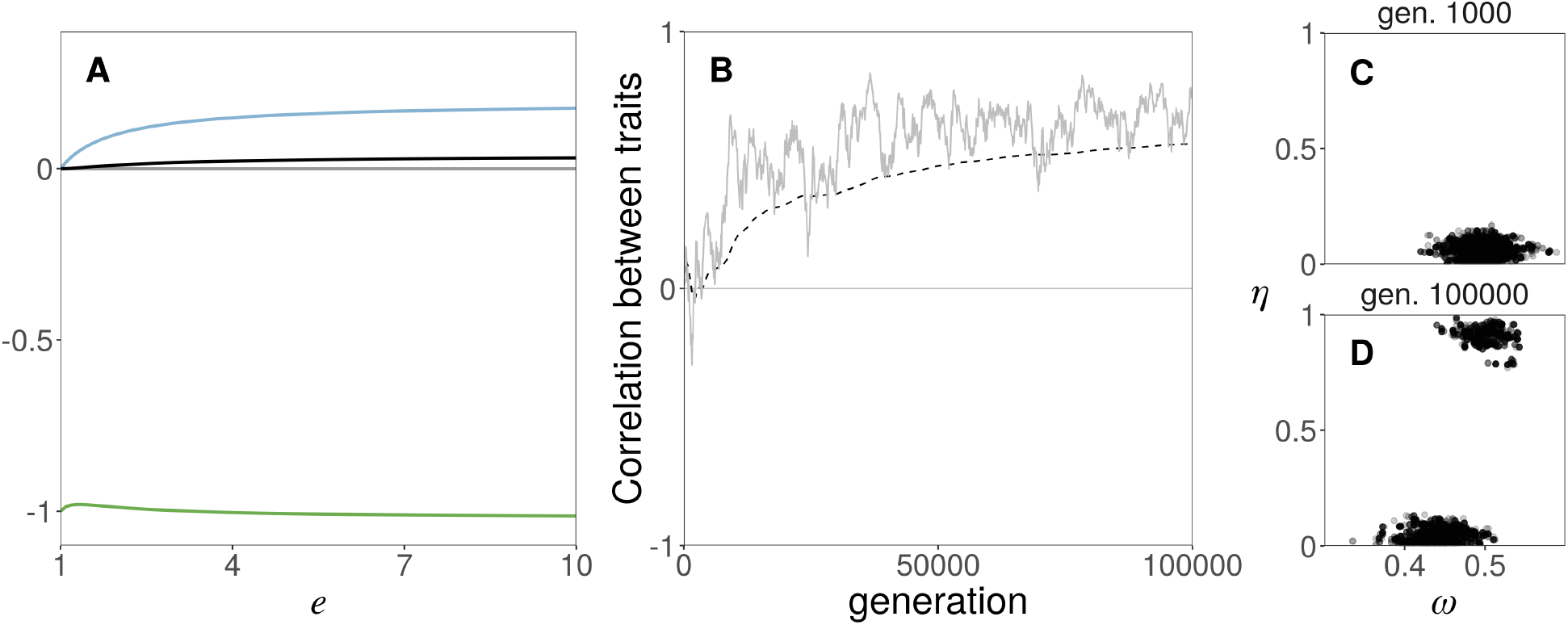
Polymorphism under monandry is due to positive correlational selection. **A**. Characteristics of the Hessian matrix at the internal singular strategy as a function of *e* for *m* = 1 (first column of fig. S1 for singular value): quadratic selection coefficient on *ω* (*h*_*ωω*_(***z***^∗^), in green) and on *η h*_*ηη*_(***z***^∗^), in grey); correlational selection (*h*_*ωη*_(***z***^∗^), in blue) and its relative strength (*h*_*ωη*_(***z***^∗^)^2^ − *h*_*ωω*_(***z***^∗^)*h*_*ηη*_(***z***^∗^), in black, Mathematica notebook for calculations). **B**. Correlation between genetic values of each trait within haplotypes in a simulated population (in gray, 4000 haplotypes sampled every 100 generations to compute Pearson’s correlation coefficient, same replicate as fig. 4C; cumulative mean in black dashed). **C & D** Distribution of genetic values of all haplotypes after 1000 generations (panel C) and after 100000 generations (panel D, same replicate as panel B and fig. 4C).

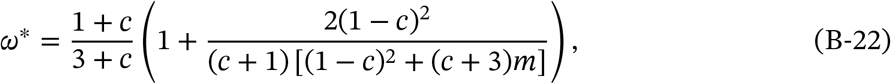

and

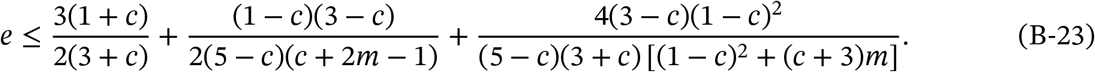

Condition eq. (B-23) corresponds to the area of the graph below the plain line in fig. 5A-B, where hybridization avoidance is stable. Conversely, the area above the plain line in fig. 5A-B (in blue) is where avoidance is not stable and thus where hybridization evolves.

###### Stability of worker-loss

Similarly, worker-loss is stable if selection on hybridization settles for an equilibrium *η*^∗^ in the absence of developmental plasticity (i.e. *s*_*η*_(***z***) = 0 for some 0 < *η*^∗^ < 1 when *ω* = 0). We find that this equilibrium reads as

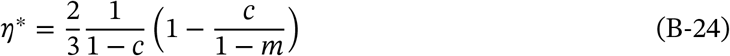

(fig. 5D). The equilibrium eq. (B-24) is between 0 and 1 (0 < *η*^∗^ < 1) and selection at this equilibrium maintains worker-loss (i.e. *s*_*ω*_(***z***) ≤ 0 when *ω* = 0 and *η* = *η*^∗^) when

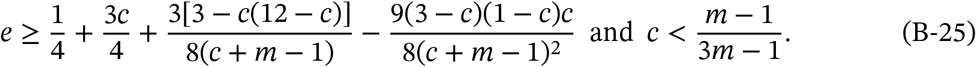

Note that condition eq. (B-25) is only possible when *m* ≥ 2. It therefore does not appear in fig. 5A (which is for the case *m* = 1) but corresponds to the area above the dotted line in fig. 5B (which has *m* = 2).

###### Worker-loss coupled with complete hybridization

In principle, it is also possible with parthenogenesis for a population to evolve worker-loss (*ω* = 0) with complete hybridization (*η* = 1) (as parthenogenesis allows the production of queens in the absence of intraspecific matings). We therefore further need to determine whether worker-loss can also be stable in the case where *η* = 1 (rather than for some 0 < *η*^∗^ < 1). We find that selection under worker-loss (*ω* = 0) and complete hybridization (*η* = 1) maintains both worker-loss and complete hybridization (i.e. *s*_*ω*_(***z***) ≤ 0 and *s*_*η*_(***z***) ≥ 0 where ***z*** = (*ω, η*) = (0, 1)) when

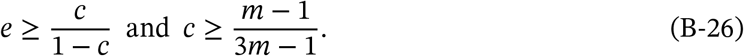

Condition eq. (B-26) corresponds to the area above the dashed line in fig. 5A-B. While condition eq. (B-25) can only be met only under polyandry (*m* > 1), condition eq. (B-26) can be met for any number of mates *m*. This means that the evolution of worker-loss under monandry and thelytokous parthenogenesis is always associated with complete hybridization in our model.

##### B.2.3 Effect of non-linear workforce productivity

Our analyses so far have assumed a linear effect of worker number on colony fitness (*α* = 1 in eq. A-2b). Here we investigate how non-linear effects of the number of workers on the pre-mating survival of virgin queens and males influence our results. We restrict our exploration to the case where queens mate with an infinite number of males and do not reproduce via parthenogenesis for simplicity (*m* → ∞ and *c* = 0). With *α* in eq (A-2b) as a variable, we find from eq. (A-8) that the selection gradient vector now reads as,

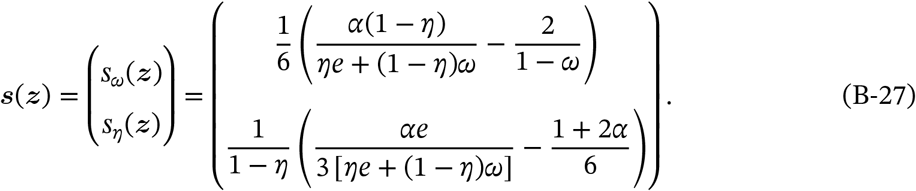

Solving for both of these gradients to vanish simultaneously, we find that there exists a unique singular strategy,

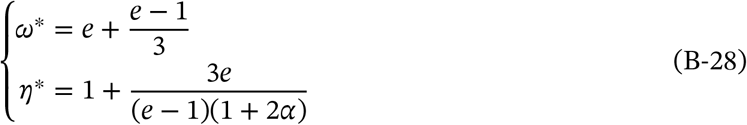

(fig. S3). The Jacobian matrix (eq. A-10) of the system eq. (B-27) at this singular value eq. (B-28) reads as

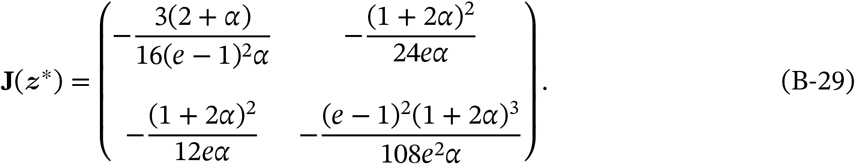

It is straightforward to show from eq. (B-29) that the singular strategy eq. (B-28) is a repellor, just as under linear effects (*α* = 1, eq. B-7). This indicates that as illustrated in fig. 3, the coevolution of caste determination and hybridization under non-linear effects also lead to either hybridization avoidance or worker-loss depending on parameters and initial conditions.

We can gain further insights into the influence of non-linear effects by determining when the singular strategy eq. (B-28) is within the trait space (i.e., when 0 < *ω*^∗^, *η*^∗^ < 1). We find that this is the case when

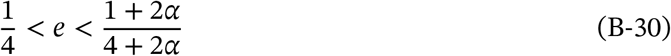

(light green region in fig. S3). This means that the threshold value for worker efficiency *e* above which worker-loss can evolve is 1/4 (as under linear effects *α* = 1). Condition (B-30) further shows that the threshold for *e* above which worker-loss always evolves (i.e. independently from initial conditions, fig. 3D for e.g.) increases with *α* (dark green region in fig. S3). In other words, the evolution of worker loss is facilitated under diminishing (*α* < 1, fig. S3A) and impaired under increasing returns (*α* > 1, fig. S3C).

**Figure S3:**
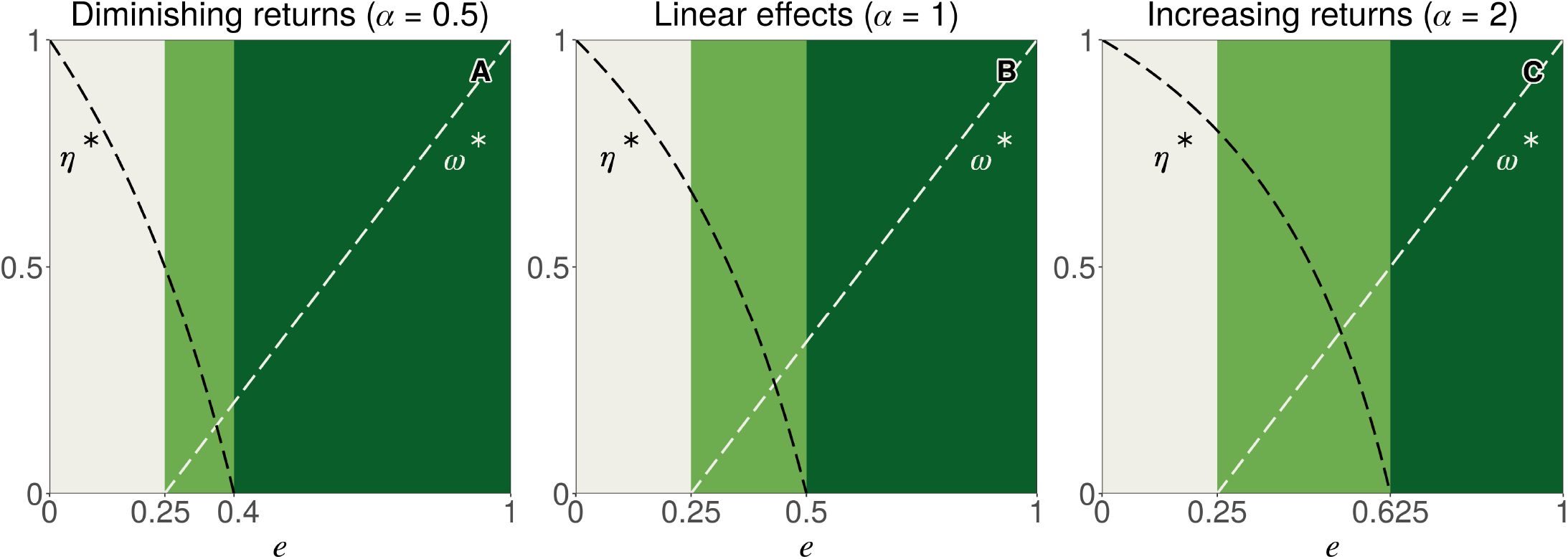
Non-linear effects of investment in workers. Singular values for *η* (in black) and *ω* (in white) as a function of hybrid worker efficiency *e* (given by eq. B-28).

